# Profiling the proximal proteome of the mu opioid receptor identifies novel regulators of receptor signaling and trafficking

**DOI:** 10.1101/2022.03.28.486115

**Authors:** Benjamin J. Polacco, Braden T. Lobingier, Emily E. Blythe, Nohely Abreu, Prachi Khare, Matthew K. Howard, Alberto J. Gonzalez-Hernandez, Jiewei Xu, Qiongyu Li, Brandon Novy, Zun Zar Chi Naing, Brian K. Shoichet, Willow Coyote-Maestas, Joshua Levitz, Nevan J. Krogan, Mark Von Zastrow, Ruth Hüttenhain

## Abstract

The mu opioid receptor (μOR), a prototypic member of the large G protein-coupled receptor (GPCR) family, represents an important target of therapeutic and abused drugs. To date, most of our understanding of μOR activity has focused on signal transducers and regulatory molecules including G proteins, GPCR kinases, and beta-arrestins. Yet it is clear that signaling through the μOR is coordinated by additional proteins recruited into the proximal interaction network of the activated receptor, which have largely remained invisible given the lack of technologies to interrogate these networks systematically.

Here, we implement a quantitative proteomics pipeline leveraging the chemical diversity of μOR agonists and APEX-based proximity labeling to investigate the protein networks that underlie μOR signaling. We leverage a novel computational framework to extract subcellular location, trafficking, and functional partners of GPCR activity from the proximity labeling datasets. Applying this unbiased, systematic approach to the μOR, we demonstrate that opioid agonists exert differences in the μOR proximal proteome mediated by endocytosis and subsequent endosomal sorting, exemplified by VPS35 and COMMD3. Moreover, we identify two novel μOR network components, EYA4 and KCTD12, that are recruited into the receptor proximal network irrespective of the activating ligand and independent of receptor trafficking but based on receptor-triggered G protein activation. We provide functional evidence that these network components form a previously unrecognized buffering system for G protein activity which broadly modulates cellular GPCR signaling.

## Introduction

The mu opioid receptor (μOR) mediates the effects of endogenously produced peptide neuromodulators and represents the main pharmacological target for a large class of exogenously administered non-peptide agonists, which are clinically used for treating pain.^1,2^ The μOR is a member of the large G protein-coupled receptor (GPCR) superfamily which operates essentially by allostery, coupling agonist-induced effects on receptor conformation to the binding of cytoplasmic proteins which mediate downstream signaling and regulation.^3^ While the interactions of the μOR with known proteins such as G proteins, GPCR kinases (GRKs) and arrestins are well-characterized, it is clear that signaling through the μOR is coordinated by additional proteins in the proximal receptor interaction network operating as signal transducers, organizers, or regulators.^4,5^ Moreover, little is known how the interactions engaged by the receptor are organized and regulated in time and space in intact cells. The discovery of novel μOR proximal interactors and their cellular organization have largely remained invisible given the lack of technologies to interrogate these networks systematically.

Recently, we and others established GPCR-APEX, a proximity labeling method based on fusing an engineered ascorbic acid peroxidase (APEX) to the receptor^6,7^, taking snapshots of the proximal receptor proteome with minute resolution, and analyzing the receptor-proximal labeling profile using quantitative mass spectrometry (MS).^8,9^ We previously demonstrated that GPCR-APEX can simultaneously capture the proximal protein interaction networks and the cellular location of activated receptors in a systematic manner through the quantification of thousands of proteins that are biotinylated proximal to the receptor. However, a major limitation to the application of GPCR-APEX has been to resolve from the complex proteomic datasets information about changes in the local interaction network from changes in the subcellular location of receptors.^8^

Here, we describe a novel computational framework which (1) models time dependent subcellular location of the activated receptor by utilizing a system of spatially specific APEX references and (2) quantitatively deconvolutes the effect of receptor location and proximal interactors in the proximity labeling data. We apply this framework to the μOR after activation with three chemically diverse agonists – the opioid peptide agonist DAMGO, the opiate alkaloid partial agonist morphine, and the chemically distinct ‘biased’ partial agonist PZM21.^10^ We reveal significantly different effects on the μOR-proximal protein environment across the agonists. We establish that the majority of these differences are driven by agonist-selective receptor trafficking in the endocytic pathway and that these, in turn, are associated with agonist-selective differences in μOR recruitment of endogenous β-arrestins. Moreover, we show that GPCR-APEX also has the ability to discover novel μOR proximal network components modulating receptor trafficking and signaling. VPS35 and COMMD3, two proteins primarily labeled in the proximity of the DAMGO-activated μOR, influence receptor distribution between cell surface and intracellular compartments. Furthermore, we discovered that EYA4 and KCTD12, which are recruited into the μOR-proximal network by all of the agonists tested and are dependent on G_i_ activity, have significant, but distinct effects on μOR signaling by interacting directly with the G protein subunits rather than the receptor itself.

## Results

### Proximal proteome changes of the activated ***μ***OR are driven primarily by differences in subcellular location

We first sought to use our approach combining APEX-based proximity biotin labeling and quantitative MS to investigate, in an unbiased manner, how activation of the μOR controls the proximal proteome environment.^8^ To comprehensively map proximal proteome changes of the μOR, we chose three agonists, DAMGO, morphine, and PZM21^10^, representing distinct chemistries as well as different efficacies and cellular responses to receptor activation. DAMGO is a peptide full agonist which causes strong activation of both G protein and β-arrestin pathways.^11^ Morphine represents a naturally occurring alkaloid and partial agonist for both G protein and β-arrestin pathways.^11,12^ PZM21 is a synthetic μOR agonist unrelated to classic morphinans, which exhibits partial agonism for G protein activation and negligible activation of the β-arrestin pathway.^10^ First, we stably expressed the μOR-APEX fusion construct in HEK293 cells, which retained receptor function with regard to signaling, internalization, and recycling (**Extended Data Fig. 1A-C**). To perform proximity labeling, the cells were pretreated with biotin-phenol followed by activation of the μOR using DAMGO, morphine, or PZM21 at a concentration of 10 µM over a time course of up to 60 minutes (**Figure 1A**). We used saturating concentrations for all ligands to ensure full receptor occupancy and therefore maximal receptor activation and biotinylation of proximal proteins. At selected time points after receptor activation, we initiated proximal biotin labeling by addition of hydrogen peroxide (H_2_O_2_) for 30 seconds, followed by quenching of the reaction, cell lysis, and enrichment of biotinylated proteins using streptavidin. To quantify relative abundance changes of biotin-labeled proteins after agonist stimulation, we utilized a combination of quantitative proteomics approaches encompassing unbiased proteomics as well as targeted proteomics based on selected reaction monitoring (SRM).

**Figure 1.**
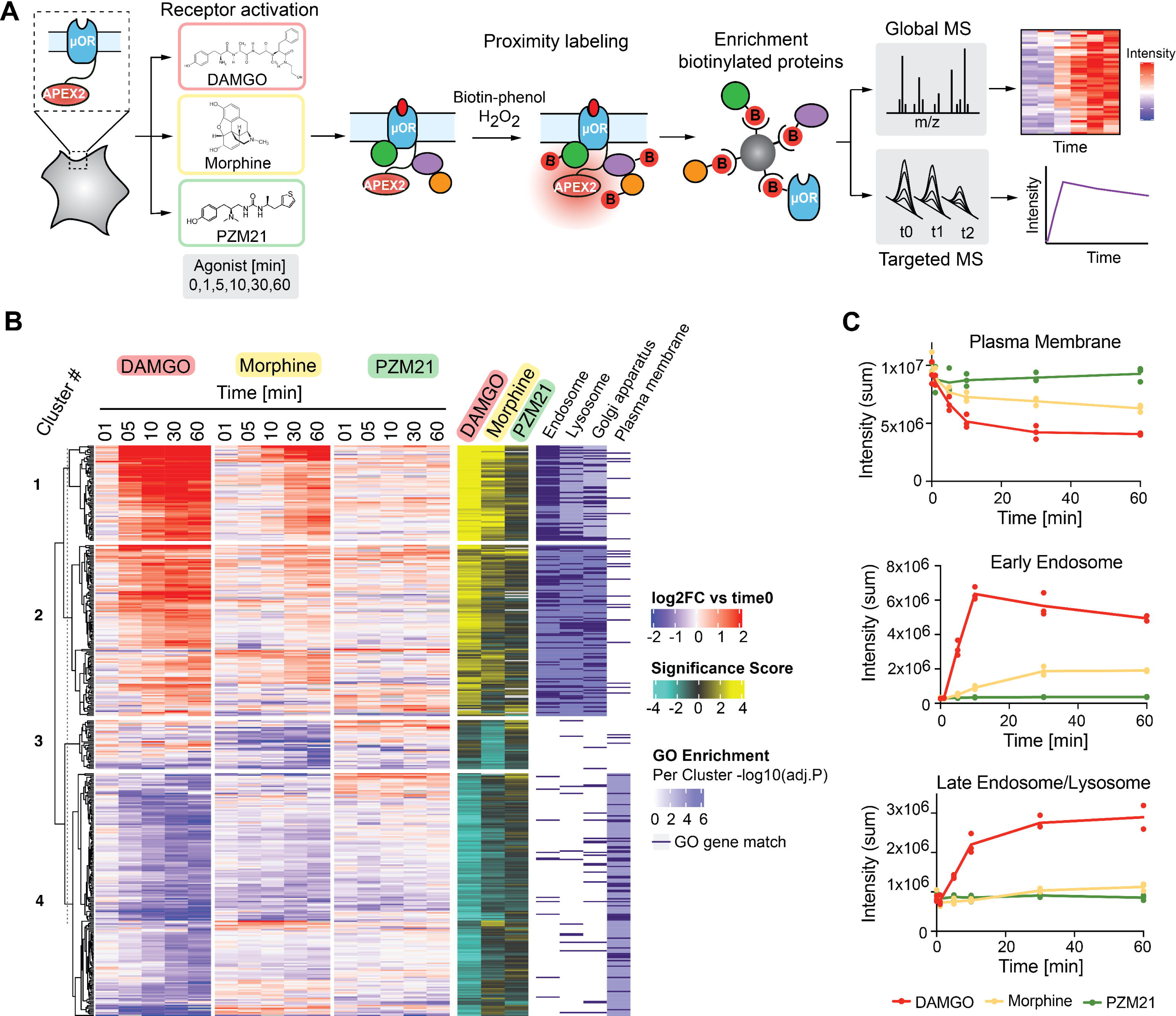
Ligand-dependent proximal proteome changes of the mu-opioid receptor are driven by cellular location. A. Experimental design of the study. μOR-APEX constructs were stably expressed in HEK cells. The receptor was activated with 10µM DAMGO, morphine and PZM21 over a time course of 60 min. Cells were pretreated with biotinphenol for 30 min and biotinylation was initiated by the addition of H_2_O_2_ for 30 sec at indicated time points after agonist treatment. Following cell lysis, biotinylated proteins were enriched using streptavidin and subsequently quantified using global and targeted mass spectrometry approaches. B. Global agonist-dependent changes in proximal proteome of the μOR. Heatmap visualizing all proteins with a significant (p < 0.01, log_2_ fold change > 1) change in biotin labeling for at least one of the ligands across the time course. The heatmap was clustered according to the significance score, calculated as a combination of the -log_10_ p-value and log_2_ fold change. Gene ontology enrichment analysis was performed for individual clusters and significant gene ontology terms for the individual clusters as well as the matching genes are indicated in dark purple. Data from three biological replicates are presented as mean. C. Targeted proteomics analysis of agonist-dependent biotinylation by μOR-APEX2 of selected localization markers for plasma membrane (top), early endosome (middle), and late endosome/lysosome (bottom). Data from three biological replicates for DAMGO (red), morphine (yellow), and PZM21 (green) are presented.

We leveraged the hundreds of proteins identified in our proximity labeling data to determine how μOR trafficking, and thus the subcellular location of the receptor, changed in a time-and-agonist dependent manner. (**Figure 1B**). To this end, we performed a statistical analysis determining proteins with significant changes in biotin labeling over the time course. For each observed protein and ligand, we scored changes in biotin labeling by fitting the time course with a polynomial curve and performed an F-test comparing it against a model with no time term. We then calculated a single significance score for each protein and ligand that summarizes the confidence, strength, and direction of the biotin labeling changes by combining the maximum fold change over the time course as a measure of strength and direction with the p-value as a measure for confidence (see **Methods** and **Supplementary Table 1**). Following hierarchical clustering of the significant proteins based on their significance score across the three ligands, we performed gene ontology (GO) enrichment analysis for each cluster.

This unbiased view of the proximal protein environment of the activated μOR indicated that (1) the majority of changes in biotin-labeled proteins are evoked by receptor endocytosis and trafficking and (2) the three ligands differ strongly in their capacity to induce endocytosis and trafficking (**Figure 1B, Extended Data Fig. 1D**). To assess ligand-dependent receptor localization quantitatively, selected localization markers for plasma membrane, early endosome, and late endosome/lysosome were monitored using targeted proteomics (**Supplementary Table 2**). We observed that DAMGO shifted the distribution of the μOR from the plasma membrane to endosomes and lysosome, while morphine resulted in less endocytosis and, astonishingly, PZM21 led to no detectable changes in μOR localization at the plasma membrane at all, something perhaps consistent with the functional selectivity for which it was developed^10^ (**Figure 1C**). Fluorescent imaging confirmed co-localization of the DAMGO-activated μOR with the early endosomal marker EEA1, but little and negligible co-localization upon activation with morphine and PZM21, respectively (**Extended Data Fig. 1E**). Together these data underscore how structurally distinct ligands can strongly influence the subcellular location of a receptor and highlight the need for a systematic approach to identify the location-specific proximal proteome(s).

### Computational framework to model receptor trafficking

To quantitatively model μOR location across multiple subcellular compartments we used our recently introduced system of spatially specific APEX-references, i.e. APEX constructs that are localized to selected subcellular compartments.^8^ Previously, we gathered information from independent trafficking assays to determine how to combine the reference conditions to reflect the spatial position of a GPCR at a particular time point after agonist addition. Here we wanted to develop a solution that would extract that information from the proximity labeling datasets themselves, thus requiring no a priori knowledge of receptor trafficking. Therefore, we developed a computational framework to quantitatively determine the localization of the spatial controls for each ligand and time point and thus more precisely deconvolve the complex proteomic profiles captured by µOR-APEX, which can traffic to different compartments over longer timescales. The aim of this comprehensive analysis is to accurately discern (1) spatial bystanders, i.e. location-specific proteins that reside in the local environment of the receptor but do not physically interact or directly participate in its function and (2) proximal protein networks that can regulate receptor signaling and trafficking.

First, we selected relevant spatial references for the plasma membrane (PM-APEX), early endosome (Endo-APEX), and lysosome (Lyso-APEX) (**Figure 2A**), which were validated to localize to and biotinylate proteins at the respective subcellular compartment (**Extended Data Fig. 2A, B**). The spatial reference samples were collected and analyzed in parallel with the μOR proximity labeling samples. We then determined indicative proteins for the subcellular locations by pairwise comparison of the biotin labeling data from the three spatial references using statistical models implemented in MSstats.^13^ Proteins were considered location indicators if they (1) showed a p-value below 0.005 and a log2 fold change (Log2FC) higher than 1.0 for at least one of the comparisons and (2) were consistently quantified across all replicates above a defined intensity threshold (**Figure 2B, Supplementary Table 3**). Next, we used these indicator proteins to quantitatively estimate the ligand-dependent fraction of the μOR at a given cellular location across the time course. Specifically, we calculated location coefficients for each ligand, time point, and biological replicate by solving a linear model based on the protein intensities of the location indicators across the spatial references and across the μOR APEX samples (**Figure 2C**). The model-based cellular location coefficients for the receptor accurately recapitulated the relative locations determined experimentally by the targeted proteomics measurements (**Figure 1C**) as well as the fluorescent imaging data (**Extended Data Fig. 1E**). Notably, our results are also consistent with previous studies investigating ligand-dependent μOR trafficking.^14^ Taken together, these results suggest that our computational framework is very powerful in modeling receptor trafficking.

**Figure 2.**
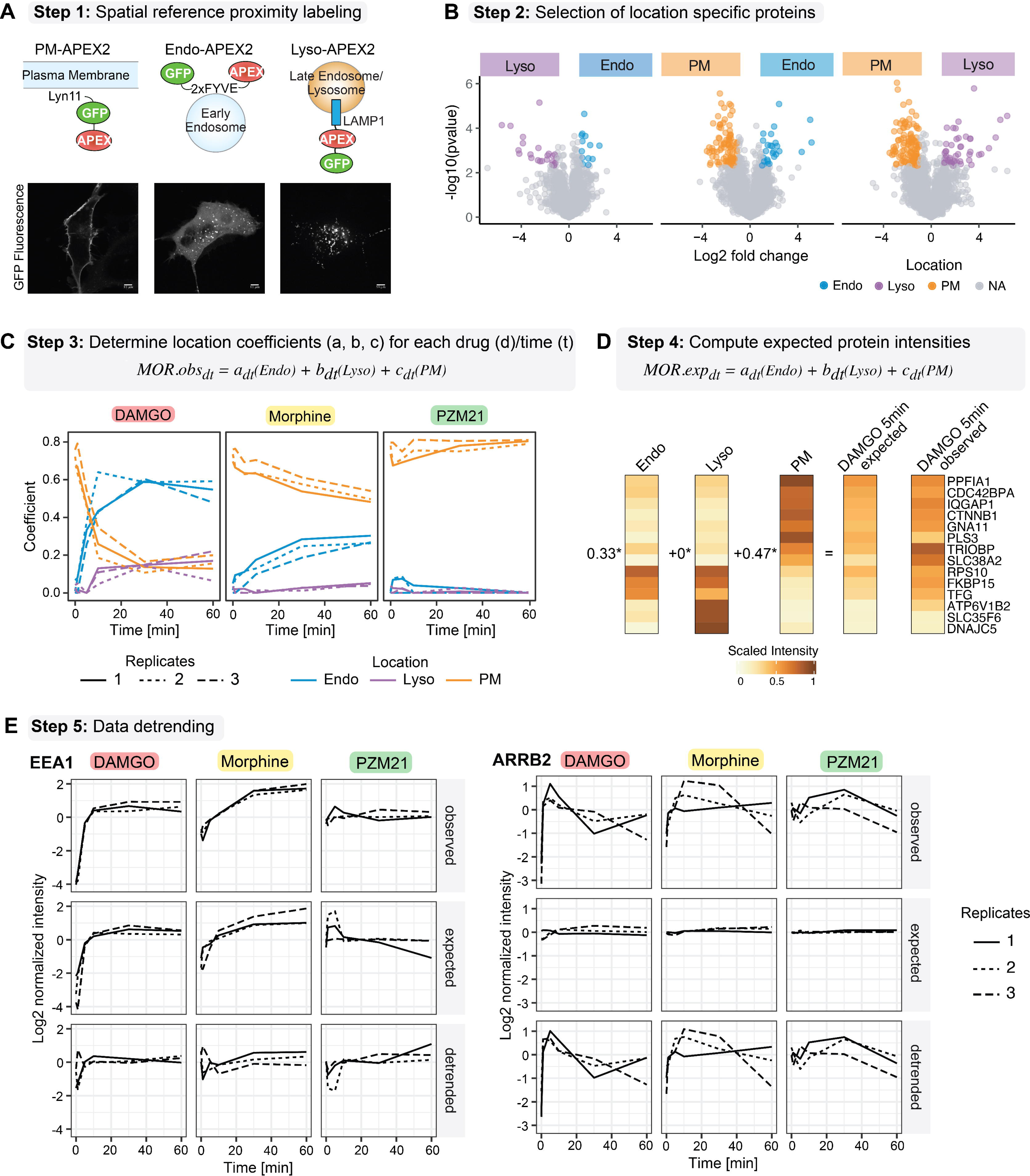
Computational framework to model ligand and time-dependent receptor location and deconvoluting receptor location from local interaction network. A. Step 1: Proximity labeling of spatial references. APEX-tagged constructs targeting APEX2 to the plasma membrane (PM-APEX2), early endosome (Endo-APEX2), or Lysosome (Lyso-APEX2) (top). Micrographs of PM-APEX2, Endo-APEX2, and Cyto-APEX2 using live-cell confocal imaging (scale bar, 10 μm) (bottom). B. Step 2: Selection of location specific indicator proteins. Volcano plots depicting pairwise comparison of spatial references. Proteins were required to (1) have a p-value below 0.005 and a fold change higher than 1.0 for at least one of the comparisons and (2) be consistently quantified across all replicates with an intensity greater than the 50th percentile to be selected as location specific proteins for plasma membrane (PM, orange), endosome (Endo, blue), and lysosome (Lyso, purple). Data from three biological replicates are presented. C. Step 3: Determination of location specific coefficients. Intensities of location specific indicator proteins are utilized to calculate coefficients for each location and ligand based on the linear model. Data from three biological replicates are presented. D. Step 4: Computing expected protein intensities. Expected intensities for all proteins quantified in the dataset are calculated based on location specific coefficients and protein intensities measured for the spatial references. A subset of the data is shown as an example. E. Step 5: Data detrending. Data detrending process to dissect localization specific effect from effect of interaction with the receptor is exemplified for EEA1 (spatial bystander) and ARRB2 (known μOR interactor). Three different temporal profiles are depicted for each protein, ligand, and replicate: the initial observed intensities (top), the expected intensities based on the location specific references (middle), and the intensities after detrending (bottom). Data from three biological replicates are presented.

We next tested whether we could leverage the model-based coefficients to deconvolve the two time-dependent profiles of spatial bystanders and proximal protein interaction networks. To this end, we calculated expected intensities for all proteins in the μOR APEX dataset based on the cellular location changes of the receptor. We combined the location coefficients for each ligand, time point, and biological replicate and the intensities of all proteins quantified across the PM, Endo, and Lyso-APEX spatial references (**Figure 2D, Extended Data Fig. 3**). Finally, based on the assumption that observed protein intensities are a combination of both μOR proximal interaction networks and spatial bystanders, we detrended the observed protein intensities for each ligand, timepoint, and replicate combination with the spatially-expected protein intensities. In brief, a protein’s expected intensity, based on spatial references and ligand/time-specific coefficients, was subtracted from its observed intensity to produce a detrended intensity (**Supplementary Table 4**). On the example of EEA1, a spatial bystander of the μOR at the early endosome and therefore an indicator for receptor trafficking, we demonstrated that detrending the data allows to account for most of the increase in biotin labeling after activation of the receptor with DAMGO and morphine (**Figure 2E**, left panel). In contrast, the agonist-dependent temporal profiles for ARRB2 (**Figure 2E**, right panel), a protein that is recruited to µOR upon activation and known to regulate receptor trafficking and signaling, as well as members of the AP2 complex (**Extended Data Fig. 4**), which directs the receptor/β-arrestin complexes into clathrin-coated pits, were not affected by the detrending.

The final outcome of the novel computational framework is the modeling of receptor trafficking directly from the proximity labeling data in contrast to measuring it independently with complementary methods as we did in our previous work. The advantages of such an approach are that it requires minimal knowledge of receptor trafficking itinerary and its kinetics, as the prediction is based on the quantitative measurement of multiple proteins. Simultaneously, the framework can subtract location-specific trends (such as EEA1) from the µOR-APEX dataset to enrich functionally relevant proteins that reside in the proximal interaction network of the receptor (such as ARRB2 and AP2 complex members). Importantly, while we show the proof of concept for µOR here, this is readily applicable for other GPCRs and more generally signaling receptors across biology.

### Ligand-dependent proximal interaction network of activated ***μ***OR differs in proteins involved in receptor endocytosis and trafficking

After detrending the proximity labeling dataset to account for labeling of spatial bystander proteins, we sought to identify novel proteins in the proximal interaction network regulating μOR function and compare these across the three ligands. To this end we performed statistical analysis on the detrended protein intensities by fitting the time course with a polynomial curve and performing an F-test to compare it against a model with no time term. Proteins with significant changes in biotinylation (Log2FC>1 and p-value<0.001) before and after data detrending for at least one ligand were considered part of the proximal interaction network of the μOR (47 proteins) and were clustered based on their significance score across the three ligands (**Figure 3A, Supplementary Table 5**). We applied a more stringent cutoff significance for the detrended dataset to focus on the proteins with higher confidence in proximity labeling changes.

**Figure 3.**
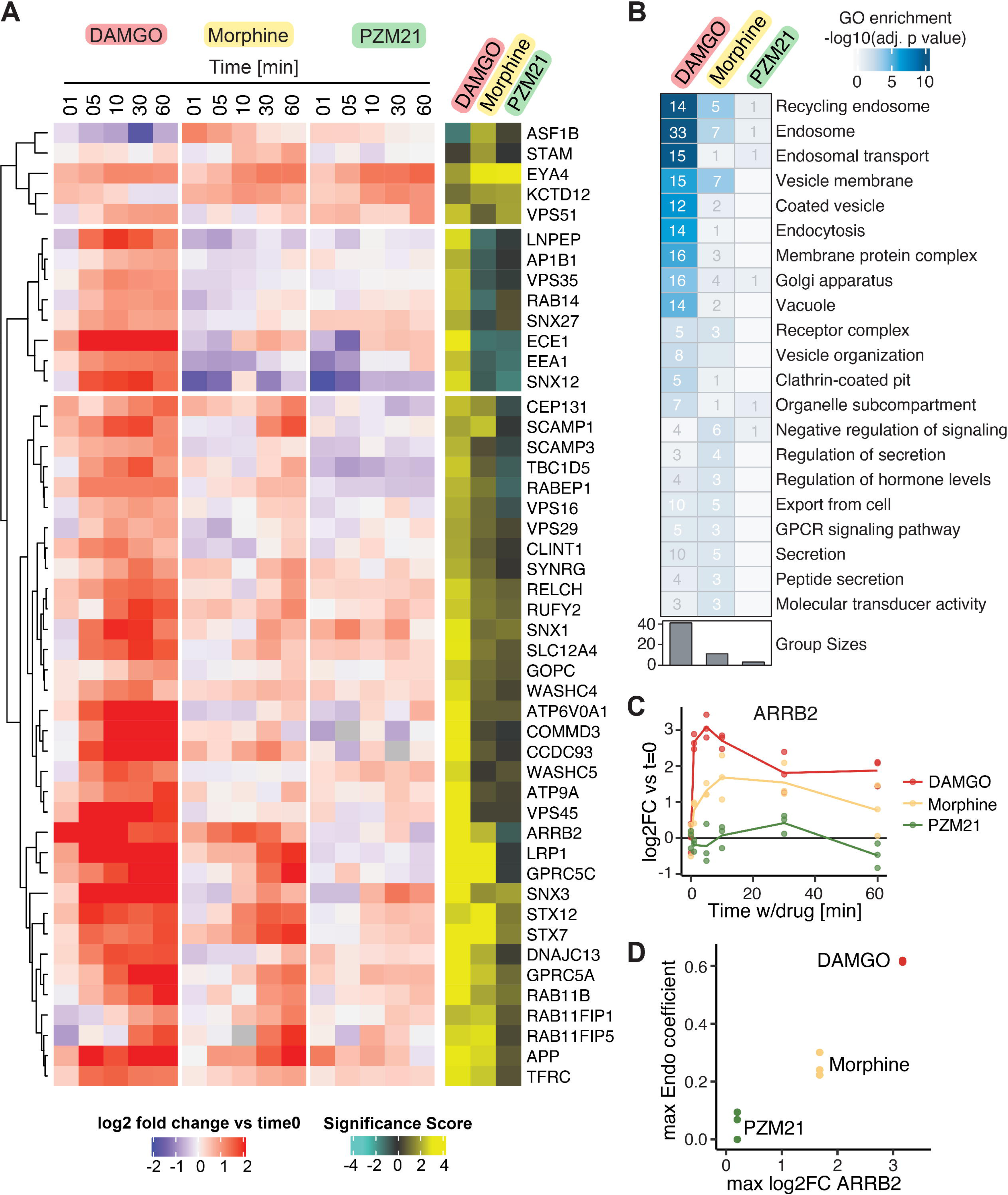
Ligand-dependent proximal interaction network of *μ*OR differs in interactors involved in receptor endocytosis and trafficking. A. Ligand-dependent proximal protein interaction networks of μOR. All proteins are visualized in the heatmap that showed a significant difference in biotin labeling (log2FC > 1 and p-value < 0.001) before and after location-specific detrending for at least one of the ligands across the time course. The heatmap was clustered according to the significance score calculated as a combination of the -log_10_ p-value and log_2_ fold change. Data from three biological replicates are presented as mean. B. Gene ontology enrichment analysis for proteins of the ligand-dependent μOR proximal interaction networks. The heatmap shows all significantly enriched gene ontology terms (adjusted p-value < 0.05) among the proteins present in the proximal interaction network of the μOR including the number of proteins that match the gene ontology terms. C. Temporal profile for ARRB2. Line charts represent the log_2_ fold change over the time course of receptor activation with DAMGO (red), morphine (yellow) and PZM21 (green) after data detrending. Data from three biological replicates are presented as mean ± SEM. D. Correlation between the maximum location coefficient calculated for the Endosome (Endo) and the maximum ARRB2 log2FC over the time course of μOR activation with DAMGO (red), morphine (yellow) and PZM21 (green).

We observed that DAMGO evoked the majority of significant changes in the proximal interaction network, which are enriched for proteins regulating endocytosis and trafficking (**Figure 3B**). Conversely, the proximal interaction network of the receptor showed only a few changes when activated with PZM21 (**Figure 3A, B**). Rather than engaging completely different proteins, our data suggests that the proximal interaction networks of the μOR activated by morphine and PZM21 represent a subset of the DAMGO network likely differing in interactors involved in receptor endocytosis and trafficking (**Figure 3B**). As an example, ARRB2 showed a strong increase in biotin labeling upon activation of the μOR with DAMGO; in contrast, activation by morphine showed less recruitment of ARRB2, and activation by PZM21 did not elicit a visible change in biotin labeling of ARRB2, again consistent with its original design^10^ (**Figure 3C**). Notably, ligand-dependent changes in ARRB2 biotin labeling were correlated with the degree of μOR endocytosis as predicted by the location coefficients (**Figure 3D, Extended Data Fig. 5**), which is consistent with previous findings.^14^

As we used HEK293 cells to map the proximal interaction network of the activated μOR, which do not endogenously express the receptor, we asked whether the μOR proximal protein network derived from HEK293 cells can be recapitulated in cells expressing the receptor. Therefore, we performed a proximity labeling experiment for the DAMGO-activated μOR in the SH-SY5Y neuroblastoma cell line. The majority (40/42) of the DAMGO-evoked proximal proteome network components discovered in HEK293 cells were also captured in SH-SY5Y cells. Impressively, we observed very similar kinetics between the two systems, suggesting the conservation of fundamental machineries regulating GPCR signaling and trafficking (**Extended Data Fig. 6, Supplementary Table 6**). Furthermore, while WASHC4 or RUFY2 did not show a DAMGO-dependent response in SH-SY5Y cells, we observed a response for related family members RUFY1 and WASH2C, respectively. One exception is the hit identified in HEK293 cells, COMMD3, which showed a distinct pattern in SH-SY5Y cells. Thus, these data demonstrate that the vast majority of the proteins we identified in HEK293 cells were also found in SH-SY5Y cells, underscoring the general conservation of GPCR signaling and regulatory proteins across different cell types.

### COMMD3 and VPS35 impact subcellular distribution of the *μ*OR

Since the proximal interaction network of the DAMGO-activated μOR was enriched for proteins related to endocytosis and endosomal trafficking, we wanted to investigate whether these proteins might influence the subcellular distribution of the μOR. We first developed an assay to measure receptor distribution between cell surface and intracellular compartments using N-terminal tagging of the μOR with a HaloTag and labeling with Halo dyes (**Figure 4A**). Using this strategy we were indeed able to differentially label the cell surface and intracellular receptor pool using sequential labeling of the HaloTag first with an impermeable (JF635i) Halo dye to first saturate the cell surface receptor pool followed by a membrane permeable (JF525) Halo dye to label the intracellular pool (**Figure 4B**).

**Figure 4.**
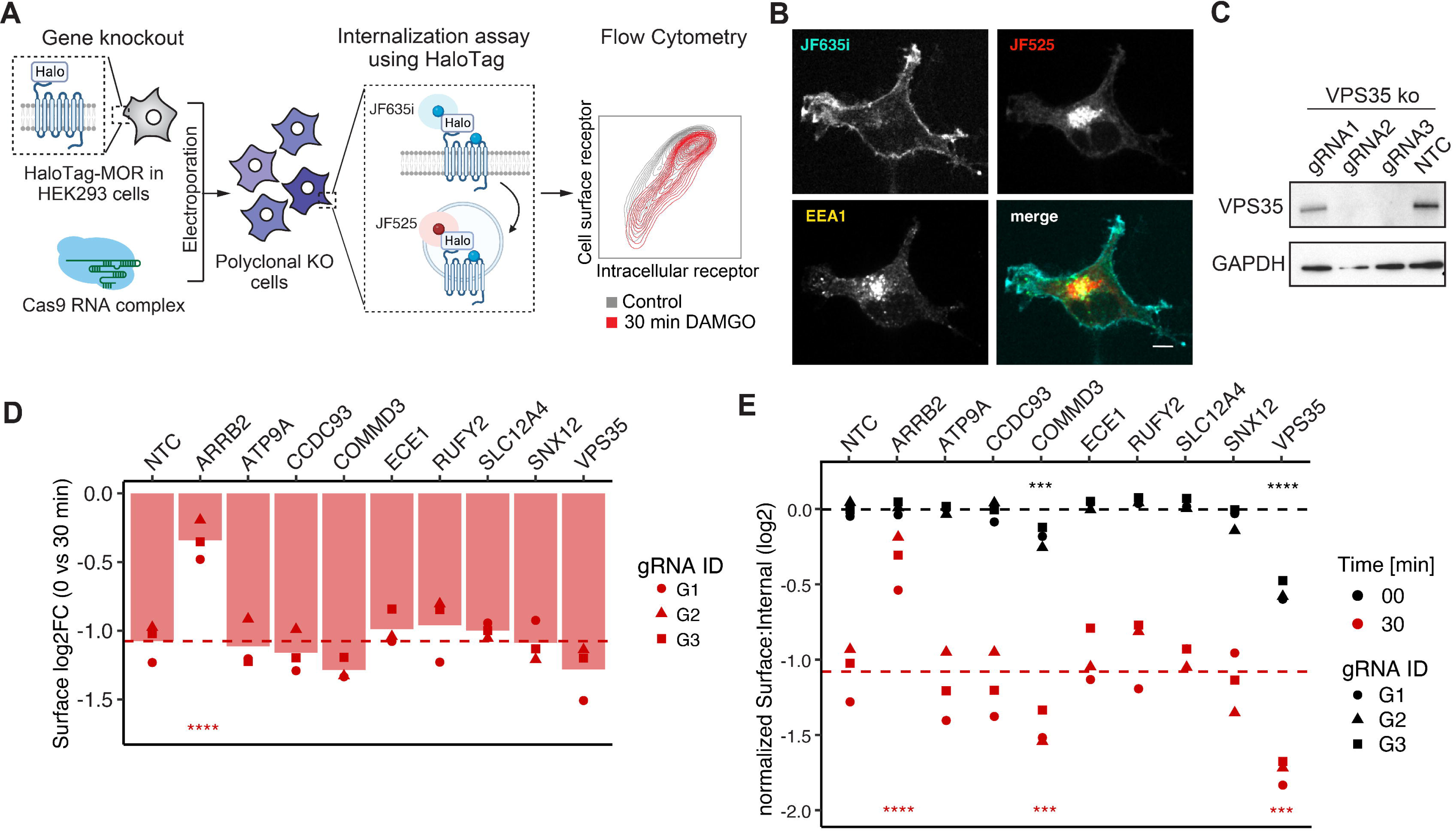
Knockout of VPS35 and COMMD3 change cellular distribution of the *μ*OR. A. Targeted CRISPR knockout screen to probe for μOR cell surface expression, internalization, and trafficking. The μOR is stably expressed in HEK293 cells with an N-terminally fused HaloTag. Polyclonal knockout cell lines are generated by nucleofection of CRISPR ribonucleoprotein complexes. Effects of gene knockout on μOR cell surface expression and internalization is assessed by receptor activation with DAMGO for 30 min followed by covalent binding of cell impermeable (JF635i) and permeable (JF525) fluorescent dyes to HaloTag to differentially label the μOR at the cell surface and intracellular compartments. B. Imaging cell surface and intracellular μOR using cell permeable and impermeable Halo dyes. JF635i was used to mark the receptor on the cell surface, JF525 was used to label the intracellular receptor. Colocalization of the intracellular μOR with endosomes was marked using an anti-EEA1. Scale bar indicates 10μm. C. Confirmation of gene knockout for three different guideRNAs targeting VPS35. GAPDH is used as loading control. D. Barplot depicts loss in cell surface μOR upon activation with 10 μM DAMGO for 30 min and in the context of CRISPR-based knockout of selected genes and non-targeting control (NTC). Line depicts the average cell surface μOR loss for the NTC. For each gene 3 guideRNAs (gRNA) were used and for each guide three biological replicates were summarized. P-value < 0.05 = *, <0.01 = **, <0.001 = ***, <0.0001 = ****. E. Plot depicts the normalized ratio comparing the population of the receptors at the cell surface and intracellularly in the presence and absence of μOR activation with 10 μM DAMGO for 30 min. Lines depicts the average normalized cell surface to intracellular ratio for NTC. For each gene 3 guideRNAs (gRNA) were used and for each guide three biological replicates were summarized. P-value < 0.05 = *, <0.01 = **, <0.001 = ***, <0.0001 = ****.

We then selected nine proximal μOR network components for CRISPR knockout to evaluate whether these functionally interact with the receptor to mediate internalization, trafficking, or recycling based on the following criteria: (1) proteins with changes in proximity labeling primarily upon receptor activation with DAMGO, (2) proteins with a proximity labeling profile that is little affected by the data detrending using our computational framework, and (3) prior evidence of the protein function in trafficking. ARRB2 was selected as a positive control expected to decrease receptor internalization, while a non-targeting control (NTC) was included as negative control. Briefly, for each gene of interest and the NTC, three guide RNAs (gRNA) were designed, complexed with Cas9 and electroporated into Halo-μOR HEK293T cells. Based on VPS35 as an example, we demonstrated high knockout efficiency by Western Blot analysis (**Figure 4C**). Effects of gene knockout on μOR cellular distribution were assessed before and after receptor activation with DAMGO for 30 min by sequential labeling with impermeable and membrane permeable fluorescent Halo dyes and subsequent flow cytometry analysis (**Figure 4A**). For each gRNA and time point, we calculated a surface to intracellular receptor ratio and assessed DAMGO-dependent receptor redistribution by comparing the surface to intracellular receptor ratios before and after μOR activation (**Figure 4D**). As expected, ARRB2 knockout resulted in significantly reduced receptor internalization compared to the NTC (**Figure 4D**). Intriguingly, looking at the surface to intracellular receptor ratios individually at each time point, ARRB2 increased the ratio after DAMGO treatment, however in contrast VPS35 and COMMD3 KO decreased the surface to intracellular receptor ratio with and without μOR activation (**Figure 4E**). Thus VPS35 and COMMD3 are recruited into the MOR interaction network selectively by DAMGO, similar to ARRB2, but they produce functional effects on MOR trafficking that are distinct from ARRB2. Together, these results support the ability of our methodology to identify novel agonist-specific network components with different effects relative to previously known network components.

### EYA4 and KCTD12 are novel G protein-dependent factors recruited in the proximal interaction network of the ***μ***OR

Despite the observed ligand-dependent differences in the μOR proximal interaction network, we discovered two proteins, EYA4 and KCTD12, showing an increase in biotin labeling upon receptor activation with all ligands (**Figure 5A**). Notably, while EYA4 shows a similar profile across all ligands, the temporal profile for KCTD12 upon activation of the μOR with DAMGO differs compared to PZM21 and morphine. EYA4 is a multifunctional protein, which functions as transcriptional co-activator when tethered to DNA through interaction with the SIX family of homeodomain proteins^15^, but also contains phosphothreonine and phosphotyrosine phosphatase activity. Interestingly, prior studies have shown that EYA2, another member of the Eyes absent (EYA) protein family, can interact and colocalize with constitutively active G_i_L proteins^16,17^ mediating an attenuation of cAMP inhibition.^17^ KCTD12 has been well characterized as an auxiliary subunit for the GABA_B_ receptor regulating the rise time and duration of G protein-coupled inwardly rectifying potassium channels (GIRKs)^18,19^ by competing with GIRK binding to the released Gβγ subunit and thus rapidly stripping Gβγ subunits from the activated channel, resulting in channel closure.^20^ Based on this prior knowledge, and the fact that EYA4 and KCTD12 were labeled by all three μOR agonists including those which stimulate negligible trafficking, we hypothesized that both proteins might regulate G protein signaling downstream of μOR. Indeed, a proximity labeling experiment of the plasma membrane (PM-APEX) upon activation of the μOR with DAMGO in the presence and absence of the G_i_□ inhibitor pertussis toxin (PTX) validated that the recruitment of KCTD12 and EYA4 to the plasma membrane is not only dependent on μOR activity, but also on G_i_ activity (**Figure 5A,B**). Interestingly, PTX treatment in the absence of μOR activation (time point 0 min) decreased biotin labeling of EYA4 at the plasma membrane significantly (log2FC -1.0, p-value 2×10^-8^, **Figure 5B**), suggesting that EYA4 might already be bound to a certain extent before μOR activation. Indeed, when we performed purifications of affinity-tagged EYA4 we detected its interaction with G_i_□ proteins at basal levels in addition to known interactors including SIX proteins (**Figure 5C**). Moreover, we found that introduction of the A633R SIX-binding mutation interfered with interaction of both G_i_□ and SIX proteins, while the D375N phosphatase-dead mutant did not alter the interactions significantly (**Figure 5C**). Finally, APEX-based proximity labeling for EYA4 indicated an increase in labeling of plasma membrane and a decrease in labeling of nuclear proteins, which was dependent on G_i_ activity (**Figure 5D,E, Extended Data Fig. 7**).

**Figure 5.**
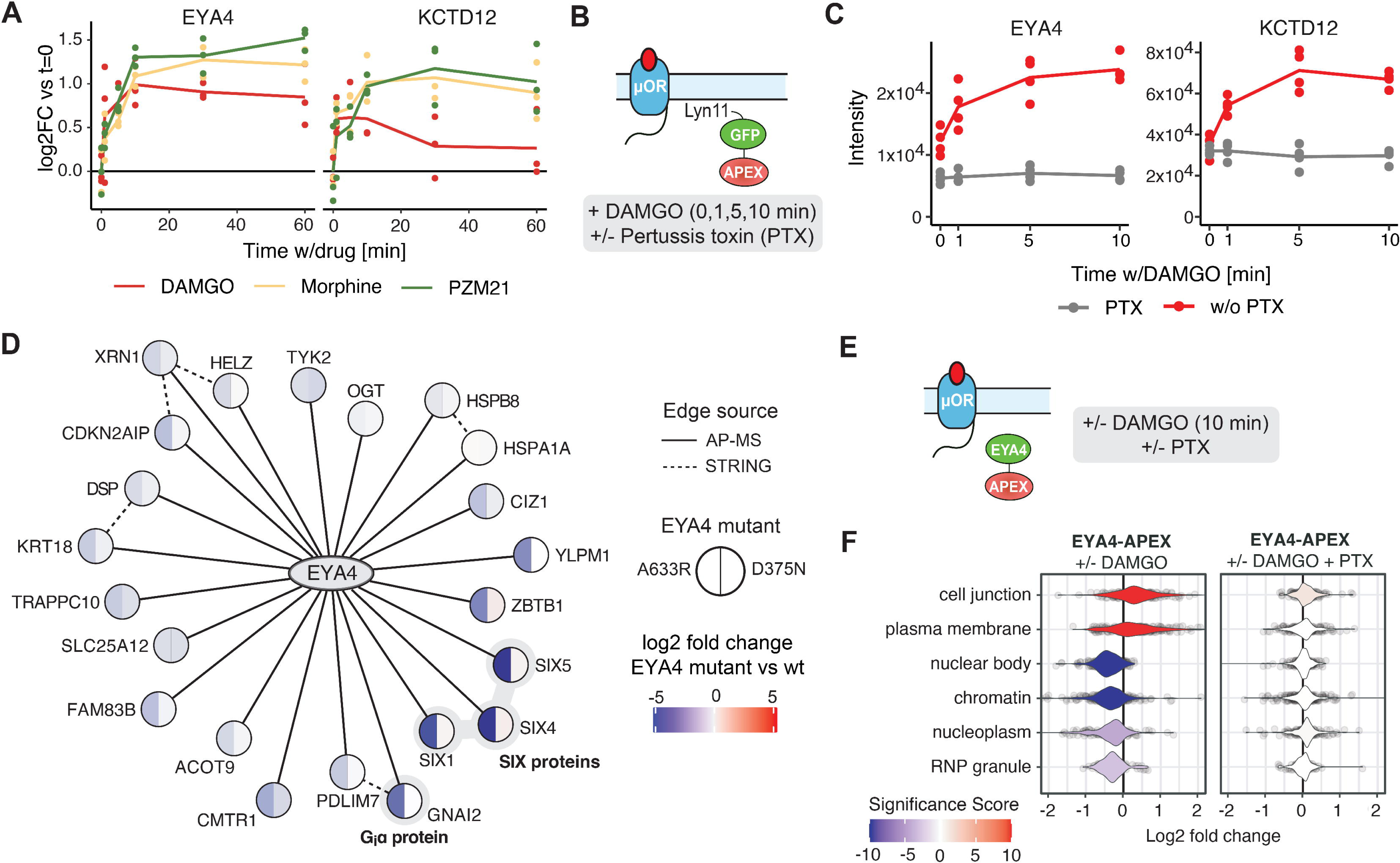
EYA4 and KCTD12 are recruited in the *μ*OR proximal interaction network in a G_i_□ activity-dependent manner. A. Temporal profile for KCTD12 and EYA4. Line charts represent the log_2_ fold change over the time course of receptor activation with DAMGO (red), morphine (yellow) and PZM21 (green) after data detrending. Data from three biological replicates are presented. B. Schematic of APEX-based proximity labeling of the plasma membrane spatial reference construct (PM-APEX) was performed in a cell line expressing the μOR in the presence and absence of the G□i inhibitor Pertussis toxin (PTX) and after activation of μOR using DAMGO. C. Line charts depict protein intensity for KCTD12 and EYA4 in PM-APEX experiment determined by quantitative MS over the DAMGO time course in the presence (grey) and absence of PTX (red). Data from four biological replicates are presented. D. Interaction network of high-confidence interactors of EYA4 determined by affinity purification. Split nodes indicate protein abundance differences for the high-confidence interactors upon introduction of SIX-binding mutant (A633R) and phosphatase-dead mutant (D375N). Data from three biological replicates are presented as mean. E. APEX-based proximity labeling of EYA4 was performed in a cell line expressing the μOR in the presence and absence of the G□i inhibitor Pertussis toxin (PTX) and after activation of the μOR using DAMGO. Volcano plot depicting log10 p-value and log2 fold change comparing biotin labeled proteins in the proximity of EYA4 in the presence and absence of μOR activation by treatment with 10 μm DAMGO for 10 min. Data from three biological replicates are presented as mean. F. Gene set enrichment analysis (GSEA) in log2FC depicted in **Extended Data Fig. 8** using the location-specific sets of proteins defined in Human Cell Map.

Taken together, our data suggest that EYA4 and KCTD12 are recruited in the proximity of the μOR in a receptor and G_i_ activity-dependent manner, and thus might function as regulators of μOR signaling through G proteins. However, based on the previously finding that the GABA_B_ seems to be the only direct receptor target of KCTD12^19^ and our experimental data for EYA4, we hypothesize that these regulatory proteins do not engage the receptor itself, but rather interact with the Gβγ and G_i_□ subunits, respectively, that are released by dissociation of the heterotrimeric G protein upon μOR activation. Thus, our data demonstrate that one of the strengths of proximity labeling is the ability to not only identify direct interactors of uOR, but also to capture and study parts of the proximal proteome which act downstream of the activated receptor.

### EYA4 and KCTD12 modulate functional signaling through G proteins

As our results indicate that EYA4 interacts with G_i_□ subunits (**Figure 5C**) and KCTD12 was previously established to bind Gβ/γ subcomplexes,^18,19^ we focused on cellular G protein signaling for initial functional assessment. We began by assaying signaling through cytoplasmic cAMP production. GPCRs stimulate or inhibit cAMP production by coupling to G_s_ and G_i_-subclass G proteins, respectively. Using isoproterenol to activate endogenous G_s_-coupled β-adrenergic receptors in HEK293 cells, we verified the characteristic G_s_-mediated cAMP response as defined by elevation of cAMP to a peak within several minutes followed by a decrease or desensitization phase in the prolonged presence of agonist (**Figure 6A left**, black curve). Using DAMGO to activate exogenously expressed μOR (blue) or somatostatin (SST) to activate endogenous G_i_-coupled somatostatin (SST) receptors (green) in HEK293 cells, we verified inhibition of the cAMP response throughout its time course and affecting the time-integrated response (**Figure 6A right**). DAMGO inhibited the response more strongly than SST, consistent with recombinant μOR being overexpressed.

**Figure 6.**
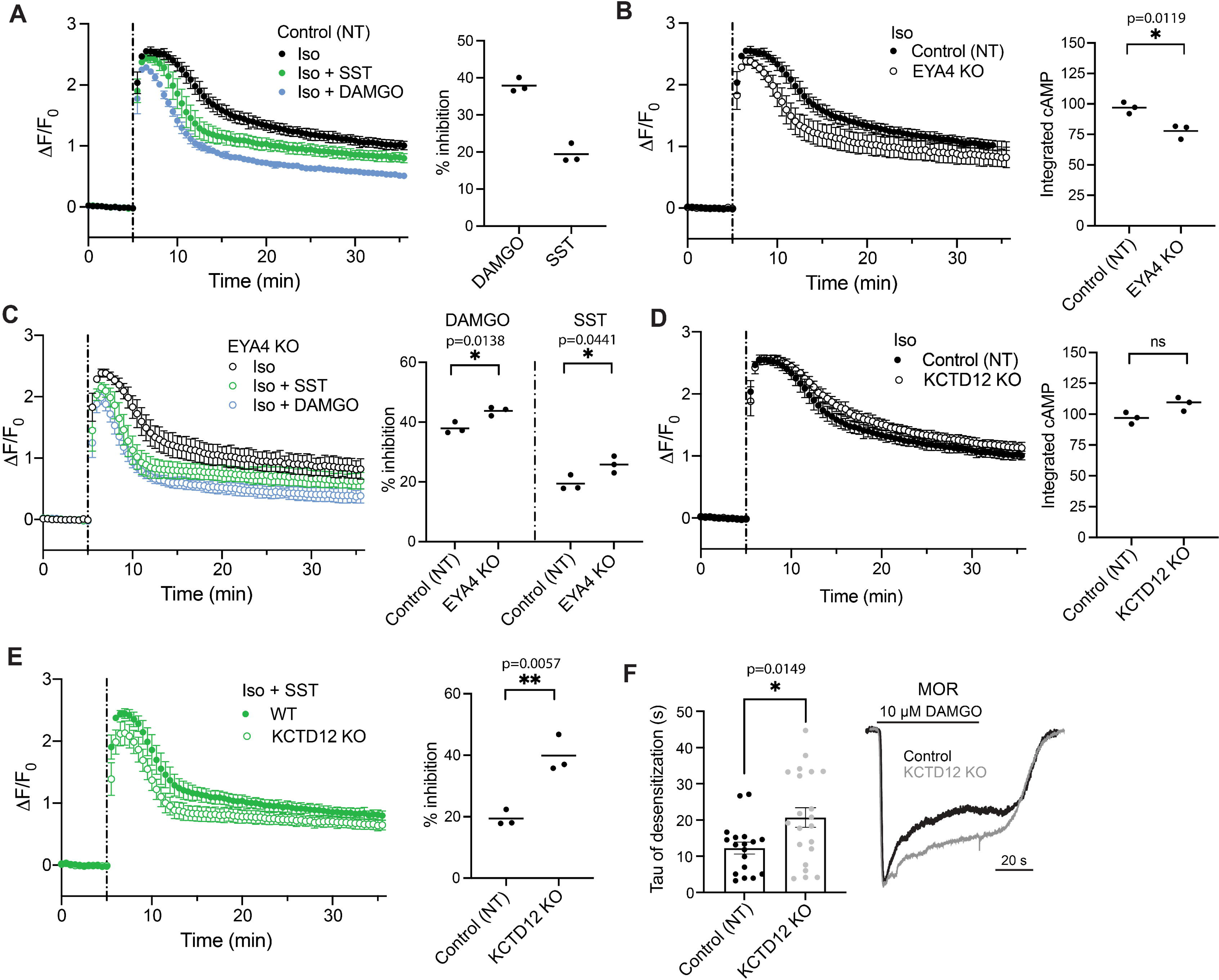
EYA4 and KCTD12 modulate G protein mediated signaling. A. cAMP activity in HEK293 non-targeting control (NT) cells stably expressing the μOR. Change in fluorescence intensity of cAMP biosensor upon stimulation with 100 nM isoproterenol (Iso) without (black) and with co-application of 1 μM somatostatin (SST, green) or 10 μM DAMGO (blue) is plotted. n = 3 B. cAMP activity in control (closed) and EYA4 KO (open) cells upon stimulation with Iso. Control curve repeated from panel A. n = 3 C. cAMP activity in EYA4 KO cells upon stimulation with Iso, with and without co-application of SST or DAMGO. Percent inhibition for Control (NT) repeated from panel A. Iso curve repeated from B. n = 3. D. cAMP activity in control (closed) and KCTD12 KO (open) HEK293 cells stably expressing the μOR upon stimulation with Iso. Control curve repeated from panel A. n = 3 E. cAMP activity in control and KCTD12 KO cells upon stimulation with Iso and SST. Percent inhibition for control repeated from panel A. n = 3. For all cAMP panels, data represent biological replicates, shown as individual data points or mean ± s.d., and significance was determined by unpaired t-test. F. Left, Quantification of the tau of desensitization of μOR-mediated GIRK currents over 60 s 10 µM DAMGO application, in control and KCTD12 KO HEK cells. Each point represents an individual cell. Unpaired t-test, *p=0.0149. Error bars represent SEM. Right, representative whole cell patch clamp recording displaying GIRK currents mediated by μOR activation over 60 s DAMGO, in control and KCTD12 KO cells.

We then applied the cAMP assay to investigate effects of EYA4 and KCTD12 on both Gs- and Gi-mediated signaling. Depleting endogenous EYA4 by CRISPR-mediated gene knockout (EYA4 KO, **Extended Data Fig. 8**) suppressed the isoproterenol-induced cAMP response over its full time course (**Figure 6B left**, open circles), resulting in a significant reduction of the integrated cAMP response (**Figure 6B right**). A similar effect was observed in an independent KO cell clone (**Extended Data Fig. 9A,B**) as well as in a polyclonal population of cells where EYA4 was partially knocked out using CRISPR (**Extended Data Fig. 9C,D**). Notably, the magnitude of inhibition produced by EYA4 KO was comparable to that produced by SST in control (non-targeting, or NT) cells (compare **Figure 6A,B**), suggesting that this effect is in a functionally relevant range. However, EYA4 KO did not prevent the effect of Gi because both SST and DAMGO were still able to further suppress the Iso-induced cAMP response in EYA4 KO cells and do so to a higher degree than in NT cells (**Figure 6C, Extended Data Fig. 9C,D**). The overall effects of EYA4 on G protein signaling cannot be fully explained by modulation of Gi alone, as PTX treatment did restore the effect on Gi-mediated cAMP inhibition (**Extended Data Fig 9E, left**), but did not restore Gs-mediated cAMP production to NT levels in EYA4 KO cells (**Extended Data Fig 9E, right**).

KCTD12 KO did not detectably change the peak cAMP response but it produced an apparent slowing of desensitization. This was a small effect, however, which did not result in a significant change in the integrated cAMP response (**Figure 6D**). A similar but more pronounced effect was observed in the second KCTD12 KO clone, resulting in a significant increase of the integrated isoproterenol-induced cAMP response (**Extended Data Fig. 9F**). KCTD12 KO also appeared to enhance G_i_-mediated inhibition of the cAMP response through endogenous SST receptors, and this effect was significant in both KO clones while varying in degree between them (**Figure 6E** and **Extended Data Fig. 9G**). We did not observe a comparable effect when G_i_-mediated inhibition was elicited through overexpressed μOR (**Extended Data Fig. 9H,I**) but were able to elicit the converse effect, reduced μOR-mediated inhibition, when KCTD12 was overexpressed together with the μOR and utilizing concentration-response analysis to assess signaling (**Extended Data Fig. 9J)**. Previous studies indicate that KCTD12 regulates signaling primarily through interactions with Gβγ subcomplexes, whereas cAMP production in HEK293 cells is regulated both by G□ and Gβγ subunits. Thus we considered that the cAMP assay may not be an ideal readout of KCTD12 effects. Accordingly, we tested regulation of overexpressed GIRK channels as a more direct readout of signaling via Gβγ. Using DAMGO to elicit Gβγ release through the μOR, we verified a rapid inward current followed by a slow 20-40% desensitization over the course of 60 seconds. KCTD12 KO did not detectably affect the initial current amplitude (**Extended Data Fig. 10A**) elicited by DAMGO but it decreased either the speed (**Figure 6F**) or extent of desensitization (**Extended Data Fig. 10B**), depending on the agonist dose applied. Conversely, overexpression of KCTD12 accelerated GIRK current desensitization (**Extended Data Fig. 10C**).

Together, these results indicate that both EYA4 and KCTD12 indeed impact cellular G protein responses, but their effects differ: EYA4 enhances the acute G_s_-mediated cAMP elevation in a manner that is consistent with it binding G_i_□, but likely involving additional homeostatic adaptation(s) of the signaling network. KCTD12 promotes signal desensitization after the peak, and it impacts both G_s_ and G_i_-mediated signaling in a manner consistent with it binding Gβγ.

## Discussion

A fundamental question for the μOR, and generally for GPCRs, is whether there are other proteins in addition to known signal transducers such as G proteins, GRKs, and Arrestins, that are recruited into the proximal interaction network of the μOR mediating the receptor’s cellular effects. To start addressing this question and to enable systematic characterization of the μOR cellular response based on receptor trafficking and its interaction networks, we advanced the previously developed GPCR-APEX methodology. We performed GPCR-APEX for the μOR after activation with three chemically diverse agonists, DAMGO, morphine, and PZM21 to determine global changes in the proximal proteome of the receptor of the time course of activation (**Figure 1**).

Previously, application of the GPCR-APEX methodology and interpretation of the datasets has been limited by (1) the data complexity resulting from receptor trafficking to multiple compartments and (2) the lack of computational approaches capable of deconvolving proximity labeling data into their constituent parts – spatial bystanders and functional proximal interaction networks. Here, we introduce a computational framework that addresses this limitation by modeling receptor trafficking based on the quantitative measurement of multiple proteins from the proximity labeling data (**Figure 2**), which requires minimal knowledge of receptor trafficking itinerary and its kinetics. This provides a significant advance to our previous work which required measurement of receptor trafficking independently with complementary methods.^8^ A prerequisite for the computational framework is the *a priori* selection of relevant spatial references either based on prior knowledge or based on information inherent in the GPCR-APEX data and their parallel processing with the GPCR-APEX samples. These spatial references allow estimating the proximity labeling background proteome followed by calculation of coefficients for each selected subcellular compartment. The proximity labeling background proteome for each compartment is determined without expression or activation of the μOR under the assumption that the majority of the proteome for each location does not change upon receptor activation. Notably, while the location coefficients do not provide an absolute measure of the receptor at a given subcellular location, they give an estimate of its relative distribution within the cell and how this distribution dynamically changes over a time course of activation. Nevertheless, we demonstrate that the coefficients provide the means to subtract the majority of location-specific trends from the GPCR-APEX datasets.

Applying this framework to the μOR, we provide a comprehensive dataset of the proximal protein environment of the receptor produced by its activation with three chemically distinct ligands – the opioid peptide agonist DAMGO, the opiate alkaloid partial agonist morphine, and the chemically distinct ‘biased’ partial agonist PZM21. These data revealed a striking ligand-dependent trafficking behavior of the μOR with subcellular resolution. Whereas DAMGO provoked strong internalization and subsequent trafficking of the μOR, this was much reduced by morphine, and was essentially undetectable by PZM21. This latter compound emerged from a structure-based effort to develop agonists functionally selective for G_i_□ versus Arrestin recruitment,^10^ leading to a compound with strong analgesic effects with reduced respiratory depression and constipation effects that are otherwise characteristic of μOR agonists. Whereas the precise mechanistic bases of the apparent functional selectivity of PZM21 and other biased agonists^21^ remains controversial,^22,23^ the differential trafficking of the μOR when activated by PZM21 does support differentiation from agonists like DAMGO and even morphine. The proteomic-based characterization of trafficking and downstream protein engagement described here offers arguably a holistic way of investigating cellular effects of novel ligands, one that may be applied to the development of other functionally selective molecules in GPCR signaling.

We then leveraged the spatial information derived from the computational framework to detrend the proximity labeling dataset for location-specific effects (**Figure 2**). Through the process of data detrending we enriched for functionally relevant proteins in the receptor interaction network (**Figure 3**), exemplified by interactors with well-characterized function in GPCR signaling and trafficking such as ARRB2 and members of the Retromer and WASH complexes. Excitingly, we also discovered novel proteins regulating μOR signaling and trafficking. Specifically, we identified COMMD3 and VPS35 as DAMGO-specific regulatory proteins of μOR cellular distribution and EYA4 and KCTD12 as common proximal interactors across all three ligands regulating receptor signaling. While our study was performed in HEK293 cells, which do not express the μOR endogenously, we were able to capture 95% of the μOR proximal interactors upon DAMGO activation in μOR expressing SH-SY5Y neuroblastoma cells (**Extended Data Fig. 6**). Impressively, we observed very similar kinetics for most proteins, including ARRB2, VPS35, EYA4 and VPS35, between the two cell systems, suggesting the conservation of fundamental machinery regulating GPCR signaling and trafficking. Nevertheless, we also observed differences in these two cell types indicating that the results obtained from HEK293 cells cannot be extrapolated to neurons without further experiments. Furthermore, the process of data detrending based on spatial references is not without limitations. It is based on the assumption that the GPCR-APEX construct and the spatial references result in similar protein labeling patterns at a given location. Given that GPCRs can traffic to multiple compartments and potentially reside in different subcellular microdomains compared to the spatial references, the simple assumption might not hold true for all proteins in the dataset. Therefore, we emphasize that functional characterization and/or validation in relevant cell systems is essential to corroborate any hypotheses derived from the proximal interaction networks.

Underscoring the ability of the GPCR-APEX/proteomics pipeline to identify novel functional interactors of GPCRs, we identify COMMD3 and VPS35 as ligand-specific regulatory proteins of μOR. We showed that knockout of COMMD3 and VPS35, two proximal μOR interactors predominantly upon activation with DAMGO, causes a shift in cell surface to intracellular distribution of the receptor (**Figure 4**). These observations are consistent with COMMD3 and VPS35 acting at a post-endocytic step of μOR trafficking. VPS35 is a subunit of the endosomal sorting complex Retromer and COMMD3 is a subunit of the Commander sorting complex.^24^ The Retromer and Commander complexes mediate endosomal sorting and recycling of a largely non-overlapping set of membrane proteins. Retromer binds membrane proteins with Ωx[L/M/V] (where Ω is any aromatic amino acid) or, when in complex with SNX27, [-][-]x[-][S/T]xL-COOH motifs (where - is any negatively charged amino acid).^25^ For example, SNX27/Retromer has been shown to mediate endosomal recycling of multiple GPCRs including B2AR and PTH1R.^26–29^ Conversely, the Commander complex consists of two sub-complexes (CCC and Retriever) and, via Retriever, bind membrane proteins with an NPxY motif.^30,31^ Intriguingly, the recycling motif in μOR, LENLE, does not match the motifs bound by either Retromer or Retriever. Thus, our results add to the emerging view that multiple sorting interactions occur in the endosomal limiting membrane and further studies will be required to determine how VPS35 and COMMD3 function in endosomal trafficking of μOR.

We also discovered components in the proximal interaction network of the μOR, EYA4 and KCTD12, that are recruited upon μOR activation by all ligands and in a G_i_-dependent manner (**Figure 5**). Our results suggest that these proximal network components are functionally relevant and that each produces distinct effects on cellular signaling by GPCRs. For EYA4, our KO data suggest that endogenous EYA4 both 1) enhances Gs-mediated stimulation of cytoplasmic cAMP elevation and 2) limits the ability of Gi to suppress this response. The second effect may result from EYA4 binding preferentially to activated G_i_□, as our AP-MS data show. It is conceivable that the first effect, on Gs-mediated stimulation of cAMP, could be mediated by the same mechanism if basal Gi tone is high. We think this is not the case, however, because the Iso-induced cAMP elevation is retained in cells pre-treated with PTX, a manipulation that should prevent regulation by Gi altogether (**Extended Data Fig 9E**). Accordingly, we speculate that EYA4 has two functional signaling effects: It limits Gi signaling, likely by binding directly to activated G_i_□, and it enhances Gs-mediated stimulation through an additional effect on cellular cAMP homeostasis that remains to be delineated. While we show activity-dependent recruitment of EYA4 in the proximity of the μOR in SH-SY5Y neuroblastoma cells (**Extended Data Fig. 6**), we have not investigated the impact of EYA4 on Gi signaling in intact neurons, which would be an important future direction.

KCTD12 is a well-characterized auxiliary subunit for GABA_B_ receptors that regulates the rise time and duration of GIRK signaling by competitive binding to Gβγ subunits.^18–20^ Our study supports the hypothesis that KCTD12 can impact G protein-mediated signaling through Gβγ for other receptors, as previously suggested.^19^ In contrast to GABA_B_ receptors, however, our data indicate that KCTD12 does not constitutively bind to the C-terminal tail of the μOR. Rather, KCTD12 appears to be directly recruited to the dissociated Gβγ subunit following receptor activation. One simple model is that KCTD12 sequesters Gβγ subcomplexes produced under agonist-induced conditions, thus depleting G protein heterotrimers available for reactivation after prolonged agonist exposure. While other possibilities cannot be presently excluded, and we note recent evidence for an additional effect of KCTD12 on adenylyl cyclase activity,^32^ this simple model is attractive because it is sufficient to explain KCTD12 affecting both cAMP and GIRK signaling, as well as enhancing desensitization without changing the acute agonist response. Given that EYA4 and KCTD12 interact with dissociated G protein subunits, rather than with the receptor itself, we conclude that GPCR-APEX is sufficiently sensitive to enable the proximal interaction network engaged and regulated by GPCRs to be probed beyond the level of direct interactions with the receptor.

Overall, our proximity labeling data obtained from HEK cells does not suggest proteins that are uniquely associating with the μOR upon stimulation with morphine and PZM21, but rather suggests that the proximal interaction networks of the μOR activated by morphine and PZM21 represent a subset of the DAMGO network likely differing in interactors involved in receptor endocytosis and trafficking. However, at this point we cannot exclude that there are ligands specific interactors that were either not captured by the APEX-based proximity labeling approach or that are not expressed in HEK293T cells and might only be captured in other cell systems endogenously expressing the μOR.

An earlier study investigating ligand-dependent protein interaction networks of the μOR using APEX-based proximity labeling proteomics discovered that stimulation of the μOR with morphine, but not DAMGO, led to increased labeling of desmosomal proteins.^33^ Civciristov *et al.* proposed a model in which the proximity to desmosomes controls morphine-mediated activation of sustained ERK within the cytoplasm.^33^ While our data indicate differences in biotin labeling of desmosomal proteins comparing stimulation with DAMGO and morphine, we hypothesize that these observations rather stem from ligand dependent differences in receptor trafficking, which lead to decreased labeling of plasma membrane proteins for the DAMGO-activated μOR.

Taken together, the presented approach is broadly applicable to assess and compare ligand-specific effects on GPCR activation. To our knowledge, this is the first methodology that can simultaneously capture the multiple layers of receptor activation - location and interaction networks in an unbiased and medium throughput fashion. As such we envisage that its application will prove to be key in informing hypothesis-driven exploration of the molecular mechanisms of how chemically distinct ligands can evoke different cellular responses, but also in drug discovery efforts to characterize and prioritize novel receptor ligands for *in vivo* testing in a medium throughput manner.

## Acknowledgement

This work was supported by funding from the Defense Advanced Research Projects Agency (DARPA) under the Cooperative Agreements HR0011-19-2-0020 (to B.K.S., N.J.K., M.v.Z., and R.H.) and HR0011-20-2-0029 (to N.J.K. and R.H.). The views, opinions, and/or findings contained in this material are those of the authors and should not be interpreted as representing the official views or policies of the Department of Defense or the U.S. Government. This work further received funding from the NIH (R01DA056354 to R.H. and M.v.Z.; P01HL146366 to N.J.K. and R.H.; 1U01MH115747 to N.J.K. and M.v.Z.; R35GM124731 to J.L: R01DA010711, DA012864 and MH120212 to M.v.Z.) and an NSF Graduate Research Fellowship (to N.A). B.T.L. was a recipient of a K99/R00 (DA043607). E.B. was initially supported by an NIH/NRSA Postdoctoral Fellowship (F32CA260118) and is currently supported by a K99 (K99GM151441). M.K.H. was supported by a training grant from NIH (5T32GM139786). A.G-H. is funded by the Margarita Salas Fellowship from the Spanish Ministry of Universities. J.L. is also supported by the Rohr Family Research Scholar Award and the Irma T. Hirschl and Monique Weill-Caulier Award.

The work was carried out in the Thermo Fisher Scientific Mass Spectrometry Facility for Disease Target Discovery at the J. David Gladstone Institutes and the UCSF Center for Advanced Technology. We thank Kirsten Obernier and Manon Eckhardt for reading the manuscript and providing critical feedback and members of both the von Zastrow lab and Krogan lab for helpful advice and comments.

## Author Contributions

B.T.L., B.J.P., M.v.Z., and R.H. conceived and directed the study with input from B.K.S. and N.J.K. APEX proximity labeling: B.T.L., Q.L., and P.K. Development of data analysis workflow and APEX data analysis: B.J.P. HaloTag-based trafficking assay: M.K.H., W.C.M, and R.H. with input from E.B. and M.v.Z. Electrophysiology: N.A., A.G-H. and J.L. Construct cloning: B.T.L., J.X., and P.K. AP-MS experiments: J.X. with input from R.H. AP-MS data analysis: Z.Z.C.N. with input from R.H. Knockout cell line generation: J.X. and R.H. cAMP measurements: E.B., B.T.L., and P.K. with input from M.v.Z. Imaging: B.N., B.T.L., M.v.Z.

## Methods

Requests for resources, reagents, or questions about methods should be directed to Lead Contact Ruth Huttenhain.

### Mammalian Cell Culture Conditions

HEK293 cells were obtained from the American Type Culture Collection (ATCC, CRL-1583), cultured in Dulbecco’s modified Eagle’s medium (DMEM, GIBCO or Fisher Scientific), supplemented with 10% fetal bovine serum (UCSF Cell Culture Facility), and maintained at 37°C in a 5% CO_2_ humidified incubator. HEK293 cells stably expressing APEX2-tagged μOR (mouse) and APEX2-tagged spatial reference constructs were selected with 500 μg/mL G418 and maintained in 100 μg/mL G418. Transfections were performed using Lipofectamine 2000 (Invitrogen) or Lipofectamine 3000 (Invitrogen) for cDNA (2 uL of Lipofectamine per 1 μg of DNA). For transient DNA expression, cells were transfected 24 or 48 hr before experiments. For electrophysiology measurements, cells were seeded on poly-L-lysine-coated 18 mm coverslips. Transfections were performed using Lipofectamine 2000 (Invitrogen).

SH-SY5Y cells were obtained from the American Type Culture Collection (ATCC CRL-2266) and cultured in Dulbecco’s Modified Eagle Medium (D-MEM)/Ham’s F-12 with L-glutamine (Cytiva), supplemented with 10% fetal bovine serum (Thermo Fisher) and PenStrep (Corning), and maintained at 37°C in a 5% CO_2_ humidified incubator. APEX2-tagged μOR (human) was stable expressed using lentiviral transduction followed by selection with 2 μg/mL puromycin.

### APEX reaction, biotinylated protein enrichment and preparation for mass spectrometry analysis

For the μOR-APEX experiments the following method was used to perform biotinylation, enrichment and prepare the samples for MS analysis. 500 μM biotin-phenol was pre-incubated with cells expressing μOR-APEX2 for 30 min at 37°C. 10 μM DAMGO ([D-Ala2, N-Me-Phe4, Gly5-ol]-Enkephalin acetate salt, Sigma-Aldrich), morphine (Morphine sulfate, Sigma-Aldrich), or PZM21 (synthesized by Enamine at 98% purity as tested by NMR and LC-MS) was added for the noted period of time. For the spatial references, cells expressing PM-APEX2, Endo-APEX2, and Lyso-APEX2 were incubated with 500 μM biotin-phenol for 30 min at 37°C, no agonists were added to these cells. Immediately prior to use, H_2_O_2_ was diluted to 2 mM final in room-temperature media (DMEM+10% FBS). APEX labeling was initiated by 1:1 mixing of the H_2_O_2_ containing media (1 mM H_2_O_2_ final) with the biotin-phenol containing media at room temperature. The labeling reaction was allowed to continue for 30 s, media was removed, and the cells were washed three times in ice cold quenching buffer (TBS supplemented with 1 mM CaCl2, 10 mM sodium ascorbate, 1 mM sodium azide, and 1 mM Trolox). Cells were incubated in quenching buffer for 20 min on ice, quenching buffer was removed, and cells were lysed in RIPA (50 mM Tris, 150 mM NaCl, 1% Triton X-100, 0.5% deoxycholate, 0.1% SDS, pH 7.4) supplemented with 10 mM sodium ascorbate and protease inhibitors (Roche Complete). Samples were briefly sonicated, spun down at 10,000 x g for 10 min, the supernatant was applied to streptavidin agarose resin (Thermo), and incubated overnight at 4°C.

Streptavidin agarose resin was washed two times in RIPA buffer (50 bed volumes per wash), four times in TBS (50 bed volumes per wash), one time in 50 mM NH_4_HCO_3_, 3 M Urea (1 bed volume per wash). Samples were reduced on resin by adding TCEP (5 mM final) and incubating, with orbital shaking, for 30 min at 55°C. Samples were alkylated by adding iodoacetamide (10 mM final), covered from light and with orbital shaking, for 20 min at room temperature. The reaction was quenched upon addition of DTT (20 mM final). The streptavidin agarose resin was spun down and the buffer exchanged to 50 mM NH_4_HCO_3_, 2 M Urea. Biotinylated proteins were cleaved on resin by the incubation of trypsin overnight at 37°C (1 μg trypsin per 20 uL of streptavidin agarose). Following proteolysis, the resin was spun down by centrifugation at 1000 x g for 1 min, and supernatant collected. The resins were washed twice with 50 mM NH_4_HCO_3_, 2 M Urea and this material was pooled with the first supernatant. The sample was acidified with TFA. NEST C18 MacroSpin columns were used to desalt the peptide sample for mass spectrometric analysis.

For the PM-APEX samples, EYA4-APEX samples and μOR-APEX samples derived from SH-SY5Y cells the following method was used to perform biotinylation, enrichment and prepare the samples for MS analysis. HEK293 cells expressing the μOR (human) and either the PM-APEX or EYA4-APEX construct were incubated with 500 μM biotin-phenol at 37°C for 30 min. 10 μM DAMGO was added for 3 different time points including 1, 5, and 10 min for PM-APEX, and 10 min for EYA4-APEX. For the pertussis toxin treatment condition, cells were preincubated with 100 ng/ml pertussis toxin for 18 hr before DAMGO treatment. APEX labeling was initiated by 1:1 mixing of the H_2_O_2_ containing media (1 mM H_2_O_2_ final) with the biotin-phenol containing media at room temperature. After 45 s of the biotinylation reaction, the cells were washed three times (1 min each) with ice cold quenching buffer (PBS supplemented with 10 mM sodium ascorbate, 10 mM sodium azide, and 5 mM Trolox). Cells were then collected in 8 mL of quench buffer and pelleted by centrifugation at 4 °C for 10 min at 3000 g. For cell lysis, cells were homogenized using probe sonication in RIPA (50 mM Tris, 150 mM NaCl, 1% Triton X-100, 0.25% sodium deoxycholate, 0.25% SDS, pH 7.4) supplemented with 10 mM sodium ascorbate, 10 mM sodium azide, 5 mM Trolox, 1mM DTT, and protease inhibitors (Roche Complete). To remove the cell debris, cell lysate was centrifuged at 10,000 x g for 10 min, and the supernatant was taken for streptavidin enrichment of biotinylated proteins.

The enrichment of biotinylated proteins was automated with the KingFisher Flex (Thermo Fisher Scientific). Supernatants were incubated at 4 °C for 18 hrs with magnetic streptavidin beads (Pierce™ Streptavidin Magnetic Beads, Thermo Fisher Scientific) which were pre-washed twice with RIPA buffer. Following incubation, beads were washed three times with RIPA buffer, one time with 1 M KCl, one time with 0.1 M Na_2_CO_3_, one time with 2 M urea in 50 mM Tris-HCl (pH 8) buffer, and two times with 50 mM Tris-HCl (pH 8) buffer. Beads were maintained in 200 uL of 2 M urea in 50 mM Tris-HCl (pH 8) buffer for on-bead digestion of proteins. Samples were reduced with 5 mM TCEP at 37 °C for 30 min, followed by alkylation with 5 mM IAA at room temperature for another 30 min, which was quenched by addition of DTT (5 mM final). For tryptic digestion, 1 ug of trypsin was added to beads and incubated with shaking at 37 °C for 4 hrs. Supernatants were taken and saved for desalting using NEST C18 MicroSpin columns.

### Unbiased mass spectrometric data acquisition and protein quantification for APEX samples

Digested peptide mixtures were analyzed by LC-MS/MS on a Thermo Scientific Orbitrap Fusion tribrid mass spectrometry system equipped with a Proxeon Easy nLC 1000 ultra high-pressure liquid chromatography and autosampler system. Samples were injected onto a C18 column (25 cm x 75 μm I.D. packed with ReproSil Pur C18 AQ 1.9 μm particles) in 0.1% formic acid and then separated with an 80 min gradient from 5% to 30% ACN in 0.1% formic acid at a flow rate of 300 nl/min. The mass spectrometer collected data in a data-dependent fashion, collecting one full scan in the Orbitrap at 120,000 resolution followed by collision-induced dissociation MS/MS scans in the dual linear ion trap with a maximum cycle time of 3 seconds between each full scan. Dynamic exclusion was enabled for 30 s with a repeat count of 1. Charge state screening was employed to reject analysis of singly charged species or species for which a charge could not be assigned.

The raw data were analyzed using the MaxQuant algorithm (version 1.5.5.1) for the identification and quantification of peptides and proteins.^34^ Data were searched against the SwissProt Human database (downloaded 01/2017), augmented with the mouse Oprm and the APEX2 sequence, concatenated to a decoy database where each sequence was randomized in order to estimate the false discovery rate (FDR). Variable modifications were allowed for methionine oxidation and protein N terminus acetylation. A fixed modification was indicated for cysteine carbamidomethylation. Full trypsin specificity was required. The first search was performed with a mass accuracy of ± 20 parts per million and the main search was performed with a mass accuracy of ± 4.5 parts per million. A maximum of 5 modifications and 2 missed cleavages were allowed per peptide. The maximum charge was set to 7+. Individual peptide mass tolerances were allowed. For MS/MS matching, the mass tolerance was set to 0.8 Da and the top 8 peaks per 100 Da were analyzed. MS/MS matching was allowed for higher charge states, water and ammonia loss events. The data were filtered to obtain a peptide, protein, and site-level false discovery rate of 0.01. The minimum peptide length was 7 amino acids. Results were matched between runs with a time window of 2 min for biological replicates. Peptide-ion intensities produced by MaxQuant from the spatial references together with the time-series μOR-APEX samples were summarized to protein intensities using the R package MSstats (version 3.23.1),^13^ specifically the function *dataProcess* with default arguments except the outlier and noise filters^35^ were enabled by setting “remove_uninformative_feature_outliers=TRUE” and “featureSubset=’highQuality’”. Log-transformed protein intensities were normalized per run using an equalize-medians procedure.

Digested PM-APEX samples, EYA4-APEX samples and μOR-APEX samples derived from SH-SY5Y cells were analyzed on an Orbitrap Exploris 480 mass spectrometry system (Thermo Fisher Scientific) coupled to a Easy nLC 1200 nano-flow ultra high-pressure liquid chromatography (Thermo Fisher Scientific) interfaced via a Nanospray Flex nanoelectrospray source. Samples were reconstituted in 1% formic acid and loaded onto a C18 column (25 cm x 75 μm I.D. packed with ReproSil Pur C18 AQ 1.9 μm particles). Mobile phase A consisted of 0.1% FA, and mobile phase B consisted of 0.1% FA/80% ACN. Peptides were separated at a flow rate of 300 nl/min using a gradient increasing buffer B over 40 min to 16% B, followed by an increase over 26 min to 28% B and 4 min to 44% B. The mass spectrometer acquired data in a data-independent acquisition (DIA) mode, collecting one full scan in the Orbitrap at 120,000 resolution followed by DIA MS/MS scans within a m/z range of 350-1050 with a fragmentation window size set to 20 m/z. The resolution of orbitrap for MS2 scans was set to 15,000 and a normalized collision energy of 30 was used for fragmentation. The DIA data were analyzed with Spectronaut (Biognosys) using direct DIA analysis default parameters for the identification and quantification of proteins. Normalization in Spectronaut was turned off. Data were searched against the SwissProt Human database (downloaded 01/2017). Peptide ion intensities from Spectronaut were summarized to protein intensities using MSstats (version 3.22.1)^13^ implemented in the R package artMS (version 1.8.3) with default settings.

### Statistical analysis of unbiased MS data for APEX samples

μOR-APEX: Each protein’s trend over the time course after treatment with agonists was scored by fitting the log2 intensities with continuous polynomial curves over time using the R functions *lm* and *poly*. To better fit the rapid changes, the collected timepoints were encoded by their ranks (1, 2, 3, 4, 5, and 6 for 0, 1, 5, 10, 30, and 60 minutes). All models included an additive term for the batch–a protein’s background intensity was expected and allowed to vary between batches. Three time-dependent models were tested that varied in the maximum polynomial power that was allowed for the ranked-time model: linear, quadratic and cubic. The time-dependent models were compared with a null-model that contained only the batch term using the R function *anova* to compute an F statistic and p-value. The time model with the lowest p-value was selected and the maximum model-predicted change between time 0 and a later time was used as the maximum log2 fold change for that protein. This process was repeated separately for the three agonists.

Spatial References: Proteins that varied between spatial references were scored with a single run of the MSstats function *groupComparison* to compare between each non-redundant pair of spatial references. The input to MSstats was the entire set of spatial references with the μOR-APEX data excluded. MSstats fits a single linear mixed model for each protein with a single categorical fixed-effect term for condition (spatial reference) and a random-effect term for batch. Using this model, MSstats reports pairwise differences in means as log2 fold change (log2FC), and a pairwise p-value calculated from a t-test assuming equal variance across all spatial references. A subset of 193 location-specific proteins was selected that could reliably distinguish locations by requiring p-value < 0.005 and log2FC > 1.0 and observed intensity in all three replicates of at least one spatial reference greater than the 50th percentile of all observed protein intensities.

Spatial coefficients and spatial detrending of μOR-APEX samples: For each μOR-APEX sample, coefficients were calculated to estimate the contribution of each spatial reference to the observed protein intensity. First, protein intensities were scaled linearly between 0 and 1 by setting the maximum observed intensity (across all spatial reference and μOR-APEX samples) for each protein to 1.0, and all other observations were set to the ratio of observed/maximum for that protein. Missing values were set to zero. A matrix representing protein intensities in the spatial references for all observed proteins, **F** (3 spatial reference columns by 4291 protein rows), was constructed using mean (per spatial reference) scaled intensity. The location specific subset matrix (**S**) was defined by using only the rows of **F** that match the 193 location-specific proteins defined in the previous section. Location coefficients for each μOR-APEX sample were then calculated using the non-negative least squares procedure in the R package nnls using the location-specific matrix **S** and the vector of location-specific protein scaled intensities from each μOR-APEX sample as inputs. We found this procedure would estimate low but non-zero coefficients where they were expected to be zero (e.g. for lysosome at 0 min) likely due to fitting noise in the μOR-APEX samples that the spatial references could not account for. To minimize these effects we used modified **S** matrices that included three additional randomized columns, by sampling from **S**. We repeated this randomization and nnls procedure 100 times and used the median value for the spatial reference coefficients. The three spatial reference coefficients for each μOR-APEX sample were then combined into a matrix **C** with 3 rows (spatial references) and 54 columns (μOR-APEX samples; 3 ligands x 6 timepoints x 3 replicates). Estimates for sample-specific location components of all protein intensities were then predicted as the matrix product of **F** X **C.** Location-detrended intensity values were then calculated by log-transforming both observed (logObserved) and predicted (logPredicted) intensity values and taking the difference, *logObserved - logPredicted*.

PM-APEX: Changes in biotinylation of proteins at the plasma membrane in response to DAMGO and/or PTX were analyzed by fitting log2 protein intensity data using linear models with the R function lm. The linear models included a term for time with DAMGO as a categorical variable, a term for +/- PTX, and also the interaction term: DAMGO x PTX which measures the significance of the different response to DAMGO with-versus without-PTX.

EYA4-APEX: Changes in biotinylation of proteins neighboring EYA4 in response to DAMGO and/or PTX were analyzed by the R package artMS (version 1.8.3) which makes use of the groupComparison function with default settings implemented in MSstats (version 3.22.1).^13^

### GO Enrichment method

Sets of proteins with significant changes were tested for enrichment of Gene Ontology terms (GO Biological Process, Molecular Function and Cellular Component). The over-representation analysis (ORA) was performed using the enricher function from R package clusterProfiler (version 3.99.0).^36^ The gene ontology terms and annotations were obtained from the R annotation package org.Hs.eg.db (version 3.12.0). In order to reduce the set of significantly enriched terms (at FDR < 0.01) to a set of non-redundant GO terms, we first constructed a term tree based on distances (1-Jaccard Similarity Coefficients of shared genes in GO) between all significant terms using the R function hclust. The term tree was cut at a specific level (R function cutree, h = 0.99) to identify clusters of non-redundant gene sets. For results with multiple significant terms belonging to the same cluster, we selected the most significant term (i.e., lowest adjusted p-value).

### GSEA on changing locations of EYA4-APEX

Significant cellular location changes between conditions were measured using gene set enrichment analysis as implemented in the R package fgsea (v 1.17).^37^ Changes in protein intensity were scored using log2 fold change comparing samples and without DAMGO. Proteins were assigned to locations according to their “NMF Localization” from Human Cell Map (Version 1) downloaded on June 7, 2021 (https://humancellmap.org/resources/downloads/preys-latest.txt). The function fgsea was used with default options, and the log10 p-value with its sign changed to match direction of enrichment was used as the significance score.

### Targeted mass spectrometric data acquisition for APEX samples

SRM assays were generated for selected interactors of μOR as well as localization markers and ribosomal proteins as internal controls for normalization (**Table S7**). SRM assay generation was performed using Skyline.^38^ For all targeted proteins, proteotypic peptides and optimal transitions for identification and quantification were selected based on a spectral library generated from the shotgun MS experiments. The Skyline spectral library was used to extract optimal coordinates for the SRM assays, e.g., peptide fragments and peptide retention times. For each protein 2-5 peptides were selected based on intensity, peptide length as well as chromatographic performance. For each peptide the 3-6 best SRM transitions were selected based on intensity and peak shape.

Digested peptide mixtures were analyzed by LC-SRM on a Thermo Scientific TSQ Quantiva MS system equipped with a Proxeon Easy nLC 1200 ultra high-pressure liquid chromatography and autosampler system. Samples were injected onto a C18 column (25 cm x 75 μm I.D. packed with ReproSil Pur C18 AQ 1.9 μm particles) in 0.1% formic acid and then separated with an 80 min gradient from 5% to 40% Buffer B (90% ACN/10% water/0.1% formic acid) at a flow rate of 300 nl/min. SRM acquisition was performed operating Q1 and Q3 at 0.7 unit mass resolution. For each peptide the best 4 transitions were monitored in a scheduled fashion with a retention time window of 4 min and a cycle time fixed to 2 s. Argon was used as the collision gas at a nominal pressure of 1.5LmTorr. Collision energies were calculated by, CE = 0.0348 ∗ (m/z) + 0.4551 and CE = 0.0271 ∗ (m/z) + 1.5910 (CE, collision energy and m/z, mass to charge ratio) for doubly and triply charged precursor ions, respectively. RF lens voltages were calculated by, RF = 0.1088 ∗ (m/z) + 21.029 and RF = 0.1157 ∗ (m/z) + 0.1157 (RF, RF lens voltage and m/z, mass to charge ratio) for doubly and triply charged precursor ions, respectively. The resulting data was analyzed with Skyline for identification and quantification of peptides.^38^ MSstats (version 3.10.6) was used for statistical analysis.^13^ Normalization across samples was conducted based on selected global standard proteins (RPL18A, RPL22, RPL28, RPL30, RPL35A, RPL6, RPL9, RPL9P7, RPL9P8, RPL9P9, RPS11). Model-based sample quantification implemented in MSstats was used to calculate the intensity of each protein in each biological sample and replicate combining all SRM transition intensities. The intensities for selected localization markers for the plasma membrane, early endosome and late endosome/lysosome were summed to calculate an intensity as a measure for receptor localization to each subcellular compartment.

### EYA4 affinity purifications

For each affinity purification (EYA4 wt, EYA4 A633R mutant, EYA4 D375N, one GFP control, one empty vector control), HEK293 cells stably expressing μOR (mouse) were plated per 15-cm dish and transfected with 10 μg of individual Strep-tagged expression constructs (2 μg for Gfp) after 20–24 hours. Total plasmid was normalized to 15 μg with empty vector and complexed with PolyJet Transfection Reagent (SignaGen Laboratories) at a 1:3 μg:μl ratio of plasmid to transfection reagent based on manufacturer’s recommendations. After more than 38 hours, cells were dissociated at room temperature using Dulbecco’s Phosphate Buffered Saline without calcium and magnesium (D-PBS) supplemented with 10 mM EDTA and subsequently washed with D-PBS. Each step was followed by centrifugation at 200 × g, 4°C for 5 minutes. Cell pellets were frozen on dry ice and stored at −80°C. For each bait and control, n=3 independent biological replicates were prepared for affinity purification.

Frozen cell pellets were thawed on ice and suspended in 1 ml Lysis Buffer [IP Buffer (50 mM Tris-HCl, pH 7.4 at 4°C, 150 mM NaCl, 1 mM EDTA) supplemented with 0.5% Nonidet P 40 Substitute (NP40; Fluka Analytical) and cOmplete mini EDTA-free protease and PhosSTOP phosphatase inhibitor cocktails (Roche)]. Samples were subjected to freeze-thaw cycle before incubation on a tube rotator for 30 minutes at 4°C and centrifugation at 13,000 × g, 4°C for 15 minutes to pellet debris. Affinity purifications were performed as follows. MagStrep “type3” beads (30 μl per sample; IBA Lifesciences) were equilibrated twice with Wash Buffer (IP Buffer supplemented with 0.05% NP40) and incubated with the lysate on a tube rotator for 2 hours at 4°C. Following incubation, beads were washed twice with Wash Buffer and with IP Buffer. To directly digest bead-bound proteins, beads were resuspended in Denaturation-Reduction Buffer (2 M urea, 50 mM Tris-HCl pH 8.0, 1 mM DTT). Bead-bound proteins were denatured and reduced at 37°C for 30 min and after bringing them to room temperature, alkylated in the dark with 3 mM iodoacetamide for 45 min and quenched with 3 mM DTT for 10 min. Proteins were then incubated at 37°C, initially for 4 hours with 1.5 μl trypsin (0.5 μg/μl; Promega) and then another 1–2 hours with 0.5 μl additional trypsin. All steps were performed with constant shaking at 1,100 rpm on a ThermoMixer C incubator. Resulting peptides were combined with 50 μl 50 mM Tris-HCl, pH 8.0 used to rinse beads and acidified with trifluoroacetic acid (0.5% final, pH < 2.0). Acidified peptides were desalted for MS analysis using a BioPureSPE Mini 96-Well Plate (20mg PROTO 300 C18; The Nest Group, Inc.) according to standard protocols.

Samples were resuspended in 4% formic acid/2% acetonitrile solution and analyzed on an Q-Exactive Plus MS system (Thermo Fisher Scientific) equipped with an Easy nLC 1200 ultra-high pressure liquid chromatography system (Thermo Fisher Scientific) interfaced via a Nanospray Flex nanoelectrospray source. Samples were loaded onto a 75 μm ID C18 reverse phase column packed with 25 cm ReprosilPur 1.9 μm, 120Å particles (Dr. Maisch). Mobile phase A consisted of 0.1% FA, and mobile phase B consisted of 0.1% FA/80% ACN. Peptides were separated by an organic gradient ranging from 4.5% to 32% acetonitrile over 53 minutes, then held at 90% B for 9 minutes at a flow rate of 300 nl/min delivered by an Easy1200 nLC system (Thermo Fisher Scientific). All MS1 spectra were collected with orbitrap detection at a 70,000 resolution and a scan range from 300 to 1,500 m/z, while the 20 most abundant ions were fragmented by HCD and detected at a resolution of 17,500 in the orbitrap. All AP-MS data were searched against the SwissProt Human (downloaded 01/2017) using the default settings for MaxQuant (version 1.6.3.3).^34^ Detected peptides and proteins were filtered to 1% false discovery rate in MaxQuant, and identified proteins were then subjected to protein-protein interaction scoring with both SAINTexpress (version 3.6.3)^39^ and CompPASS.^40^ We applied a filtering strategy to determine the final list of reported interactors. All protein interactions that possess a CompPASS WD score percentile > 0.95, a SAINTexpress BFDR ≤ 0.05. CompPASS scoring was performed using a larger in-house database derived from 11 baits that were prepared and processed in an analogous manner to this AP-MS dataset. This was done to provide a more comprehensive collection of baits for comparison, to minimize the classification of non-specifically binding background proteins as high confidence interactors. To determine the mutant-dependent changes in protein interactions of EYA4, protein abundances for mutant and wildtype AP-MS were compared using MSstats (version 3.14.1).^13^ Protein interaction network was visualized using Cytoscape (version 3.8.1).^41^

### Generation and characterization of CRISPR knockout cell lines for EYA4 and KCTD12

KCTD12 and EYA4 knockout (KO) cell lines were generated by electroporation of Cas9 ribonucleoprotein complexes (Cas9 RNPs) into HEK293 cells stably expressing the μOR (mouse) followed by clonal selection and characterization of the KO. Electroporation was performed using the SF Cell Line 4D-Nucleofector X Kit S (Lonza) and 4D-Nucleofector (Lonza). Recombinant S. pyogenes Cas9 protein used in this study contains two nuclear localization signal (NLS) peptides that facilitate transport across the nuclear membrane. The protein was obtained from the QB3 Macrolab, University of California, Berkeley. Purified Cas9 protein was stored in 20 mM HEPES at pH 7.5 plus 150mM potassium chloride, 10% glycerol, and 1mM tris(2-carboxyethyl)phosphine (TCEP) at −80°C. Each crRNA and the tracrRNA was chemically synthesized (Dharmacon/Horizon) and suspended in 10 mM Tris-HCl pH 7.4 to generate 160 μM RNA stocks. To prepare Cas9 RNPs, crRNA and tracrRNA were first mixed 1:1 and incubated 30 min at 37°C to generate 80 μM crRNA:tracrRNA duplexes. An equal volume of 40 μM S. pyogenes Cas9-NLS was slowly added to the crRNA:tracrRNA and incubated for 15 min at 37°C to generate 20 μM Cas9 RNPs.

crRNAs targeting EYA4 (5′-CCGTAAGTGGGCAAGCTGTA-3′) and KCTD12 (5′-CGTGACCCGGCGCTGCACGG-3′) were designed by Dharmacon. For each reaction, roughly 3×10^5^ HEK293 cells were pelleted and suspended in 20 μL nucleofection buffer. 4 μL 20 μM Cas9 RNP mix was added directly to these cells and the entire volume transferred to the bottom of the reaction cuvette. Cells were electroporated using program CM-130 on the Amaxa 4D-Nucleofector (Lonza). 80 μL pre-warmed complete DMEM was added to each well and the cells were allowed to recover for 5 min at 37°C followed by dilution in complete DMEM media for limited dilution to generate clonal cell lines. Clonal cell lines were characterized by genomic sequencing to confirm gene editing of EYA4 and Western Blot analysis for KCTD12 (Cell Signaling, Cat# 81935S, dilution 1:1000) to assess reduction in protein expression. GAPDH (BioLegend, Cat# 607904, dilution 1:5000) was used as a loading control for the Western Blot analysis.

### Generation of polyclonal knockout cells for HaloTag assay

Stable cell lines expressing HaloTag-μOR (human) were generated using HEK293T landing pad cells. These cells contain a genetically integrated doxycycline inducible promoter, followed by a BxBI recombination site, and split iCasp9. This cell line was a gift from Doug Fowler’s group (Univ. Washington).^42^ To generate HaloTag-μOR cell lines, landing pad constructs containing the N-terminal HaloTag-μOR-P2A-PuroR fusion downstream of a BxBI-compatible *attB* site were co-transfected (1:1) with a BxBI expression construct (pCAG-NLS-Bxb1^43^) into the HEK293T landing pad cells using Lipofectamine 3000 according to the manufacturer’s instructions in a six-well dish. All cells were cultured in 1X DMEM, 10% dialyzed FBS, 1% sodium pyruvate, and 1% penicillin/streptomycin (D10) and incubated at 37°C in a 5% CO_2_ humidified incubator. 48h following transfection, expression of integrated genes or iCasp-9 selection system is induced by the addition of doxycycline (2 μg/μL) to D10 media. 48h after induction with doxycycline, AP1903 is added (10nM) to cause dimerization of Casp9. Successful BxB1 recombination shifts iCasp9 out of frame, so only non-recombined cells will die from iCasp-9 induced apoptosis following the addition of AP1903. After 48h of AP1903-Casp9 selection, the media was changed back to D10 with doxycycline and cells were allowed to recover for 48h. Cells were then frozen until use.

Polyclonal KO cells were generated by electroporation of Cas9 ribonucleoprotein complexes (Cas9 RNPs) into HEK293T cells stably expressing HaloTag-μOR (human). Electroporation was performed using the SF Cell Line 4D-Nucleofector X Kit S (Lonza) and 4D-Nucleofector (Lonza). Recombinant S. pyogenes Cas9 protein used in this study contains two nuclear localization signal (NLS) peptides that facilitate transport across the nuclear membrane. The protein was obtained from the QB3 Macrolab, University of California, Berkeley. Purified Cas9 protein was stored in 20 mM HEPES at pH 7.5 plus 150mM potassium chloride, 10% glycerol, and 1mM tris(2-carboxyethyl)phosphine (TCEP) at −80°C. Each crRNA and the tracrRNA was chemically synthesized (Dharmacon/Horizon) and suspended in 10 mM Tris-HCl pH 7.4 to generate 160 μM RNA stocks. To prepare Cas9 RNPs, crRNA and tracrRNA were first mixed 1:1 and incubated 30 min at 37°C to generate 80 μM crRNA:tracrRNA duplexes. An equal volume of 40 μM S. pyogenes Cas9-NLS was slowly added to the crRNA:tracrRNA and incubated for 15 min at 37°C to generate 20 μM Cas9 RNPs. crRNAs targeting selected proteins and non-targeting controls were designed by Dharmacon. For each reaction, roughly 2×10^5^ HEK293 cells were pelleted and suspended in 20 μL nucleofection buffer. 4 μL 20 μM Cas9 RNP mix was added directly to these cells and the entire volume transferred to the bottom of the reaction cuvette. Cells were electroporated using program CM-130 on the Amaxa 4D-Nucleofector (Lonza). 80 μL pre-warmed complete DMEM was added to each well and the cells were allowed to initially recover for 10 min at 37°C followed by diluting with DMEM media, plating into 24-well plates and further recovering for 3 days. KO for selected genes was confirmed by Western Blot analysis, e.g. for VPS35 (Abcam, Cat# ab10099, dilution 1:100) to assess reduction in protein expression. GAPDH (BioLegend, Cat# 607904, dilution 1:5000) was used as a loading control for the Western Blot analysis.

### HaloTag-based internalization assay

Following CRISPR-mediated KO of target genes, HaloTag-μOR 20k cells were seeded into each well of a poly-D-lysine coated 24-well plate in D10 media containing doxycycline (2 μg/mL). For each KO strain, there were two wells of cells plated. Following 36h of doxycycline expression of HaloTag-μOR, media was replaced with D10. For each KO line, 48h after seeding, one well is treated with 10uM DAMGO and the other is left untreated. Cells were incubated at 37C/5% CO2 during treatment with DAMGO. Following DAMGO treatment, plates were transferred to 4C for 10m to halt receptor trafficking. Subsequently, all cells were treated with a HaloTag dye, JF635i, dissolved in D10 to a final concentration of 200nM for 30 min at 4C. JF635i is a cell impermeable dye that conjugates to cell surface-expressed receptors. The JF635i dye solution was aspirated, and cells were treated with a cell permeable HaloTag dye, JF525 in D10 (no phenol red) and incubated at 4c again for 30m. The JF525 dye solution was then aspirated, and cells were washed twice with PBS. D10 (no phenol red) was then added, and cells were incubated 30m further to facilitate the washout of excess dye. Finally, cells were washed 1x with PBS-EDTA, detached using TrypLE Express for 10m at room temperature, and resuspended in PBS + 2.5% dialyzed FBS and kept on ice prior to flow analysis. Stained cells were analyzed via flow cytometry using an Attune NxT flow cytometer. 50k cells were collected for each condition. Cells were gated on FSC-A/SSC-A to separate HEK293T whole cells then FSC-A/FSC-H to find singlets. Following this, signals for each fluorescent dye, JF635i (RL1-A) and JF525 (BL1-A), were analyzed as follows.

Surface signal to interior signal ratios were normalized by multiplication by a constant factor so that the mean surface:interior ratio across all replicates and guides of the non-targeting control at time zero was equal to 1.0. Ratios above 1.0 thus have more surface signal than the NTC, and ratios below 1.0 have less surface. The following statistical procedure was then repeated three times on log2-transformed normalized ratios to find significant differences between gene knockouts and the non-targeting control for three different scores: log normalized surface:interior ratios at time 0, log normalized ratios after 30 minutes with agonist, and the difference between log normalized ratios after 30 minutes with agonist versus no agonist (aka log2FC). For the latter, the subtraction was performed on matched data points within batches. First, the data across the four different batches were summarized to a single mean value per guide by fitting a linear model (R function *lm*) with additive terms for batch and guide. Outliers were detected as data points with modeled residuals with absolute value greater than 3 standard deviations estimated on the entire model. If any outliers were discarded, the procedure was repeated, including detecting and discarding new outliers. The function *emmeans* in R package emmeans (version 1.7.3) was then used to get an average across replicates for each guide. The averages for the different guides were treated as independent measurements, and fit to a linear model (R function *lm*) with only the targeted gene as the single dependent variable, with contrast statistics (t-test) computed using *emmeans* with each gene knockout compared against the single NTC. P values were calculated using two tailed tests on the t statistics.

### Live Cell Imaging

HEK293 Cells stably expressing CMV:PM-APEX2-GFP and CMV:LYSO-APEX2-GFP were seeded onto 35 mm glass bottom (12 mm glass) imaging dishes coated with poly-L-Lysine. The cells were allowed to grow for 24 hours prior to live cell imaging. Prior to imaging the LYSO-APEX2 cells were labeled with 60 nM final Lysotracker Deep Red (Cat#: L12492) for 45 minutes at 37 C. After this labeling period the cells were washed twice with HEK293 growth media (DMEM + 10% FBS) and media was exchanged with non-phenol red containing DMEM supplemented with 10% FBS. Prior to imaging the PM-APEX2 cells were labeled with Cell Mask Deep Red (0.75x final, Cat#: C10046) for 5 minutes at room temperature to reduce internalization. After this labeling period the cells were washed following the above protocol for the LYSO-APEX2 cells. Cells were imaged using a Nikon Spinning Disk with a 100X objective (1.49 NA). A 488 nm laser line (525/38 Em filter) was used to visualize the GFP-tagged construct and a 640 nm laser line (700/75 Em filter) was used to visualize the location markers.

### Fixed Cell Imaging

HEK293 Cells stably expressing CMV:PM-APEX2-GFP, CMV:ENDO-APEX2, and CMV:LYSO-APEX2-GFP were seeded onto #1.5 thickness, 12 mm coverslips coated with poly-L-Lysine. The cells were allowed to grow for 24 hours prior to performing the APEX2-proximity labeling reaction. The cells were pre-incubated with 500 uM biotin-tyramide for 30 minutes at 37 °C in DMEM + 10% FBS. Prior to use, H2O2 was diluted to 2mM final in room temperature DMEM +10% FBS containing 20 mM HEPES. Addition of the H_2_O_2_ containing media occurred at room temperature and the reaction was allowed to proceed for 30 seconds. The reaction was then stopped on ice with a quenching buffer consisting of PBS containing 10 mM sodium ascorbate, 1mM trolox, and 10 mM sodium azide. The cells were washed with this quenching buffer three times and then fixed with 4% formaldehyde for 20 minutes at room temperature. The cells were then washed with TBS and blocked with an imaging buffer containing 3% BSA and 0.1% TritonX-100 in PBS. After an hour long blocking step the cells were labeled with an anti-APEX2 primary (1:750, Cat#: ab222414), Neutravidin-405 (1:500, Cat#: 22831), and the ENDO-APEX2 cells were labeled with an anti-EEA1 primary (1:500, Cat#: 48453S). This labeling was allowed to occur overnight at 4°C and the following day samples were washed three times with PBS and labeled with secondary antibodies (1:1000) in the same imaging buffer. Samples were then washed three times with PBS and mounted onto glass slides using Prolong Diamond Mountant (Cat#: P36961) to preserve the GFP channel for validation of the APEX2 localization. Coverslips were allowed to fix overnight and samples were imaged on a Zeiss LSM 900 with Airyscan 2, following airyscan processing in Zen Blue software (Zeiss), images were processed in FIJI.

### Flow cytometric analysis of receptor trafficking

Flow cytometric analysis of receptor surface immunofluorescence was used to determine agonist induced internalization and subsequent agonist-withdrawn surface recovery (recycling). HEK293 cells stably expressing FLAG-tagged μOR were left untreated as a control, incubated with 10 μM DAMGO for the noted time (0-30 min). To measure recycling, cells were incubated with 10 μM DAMGO for 30 min, washed, and then incubated for an additional 30 min with 10 μM naloxone. All cells were washed twice in ice-cold PBS to stop trafficking, and incubated at 4°C for 45 min with 2 μg/mL Alexa647 (Life Technologies)-conjugated M1 anti-FLAG (Sigma, Clone M1, Cat# F-3040, 1:1000 dilution). Cells were washed once in PBS at 4°C, and then mechanically lifted in PBS for an additional 45 min at 4°C. Median fluorescence intensity of 10,000 cells per condition was measured using a FACSCalibur instrument (Becton Dickinson). Internalization was calculated as a fraction of the agonist treated condition divided by untreated. Recycling was calculated as a fraction of surface recovered receptor divided by the internalized receptor. At least four independent biological experiments were performed in triplicate for each condition.

### Live cell cAMP accumulation assay

HEK293 cells stably expressing μOR or μOR-APEX2 were transiently transfected with the cAMP biosensor pGLO-20F (Promega). Prior to agonist stimulation, cells were incubated with 250 μg/mL luciferin for 45 min in DMEM without phenol red or serum. 10 μM DAMGO and 10 nM isoproterenol or 10 nM isoproterenol (reference condition) were added to each well, and placed at 37°C. Luminescence was recorded every 10 s with a CCD sensor for 20 min. Luminescence signal generated agonist stimulation was integrated across 1 minute of maximum average signal for the given condition, and normalized to the integrated signal from 1 minute of the control condition.

### cADDis cAMP biosensor assay

Real-time cAMP dynamics were measured using the Green Up cADDis cAMP biosensor (Montana Molecular) according to the manufacturer’s protocol. Briefly, HEK293 cell lines were lifted using TrypLE Express (Thermo Fisher) and resuspended in media supplemented with the appropriate volume of cADDis BacMam. Cells were plated into a 96-well plate (Corning #3340) at a concentration of 50,000 cells per well. After resting at room temperature for 30 minutes, plates were incubated under normal culture conditions overnight. Plates were washed twice with assay buffer (20 mM HEPES pH 7.4, 135 mM NaCl, 5 mM KCl, 0.4 mM MgCl_2_, 1.8 mM CaCl_2_, 5 mM d-glucose) before a ten minute incubation in a plate reader (Synergy H4, BioTek) pre-warmed to 37°C. Fluorescence was detected using an excitation wavelength of 500 nm and an emission wavelength of 530 nm every 30 seconds. After a five minute baseline reading, vehicle or agonist(s) (100 nM isoproterenol, 10 μM DAMGO, and 1 μM SST) were added, and fluorescence was measured for 30 minutes. A baseline fluorescence (F_0_) was calculated for each well by averaging its fluorescence over the five minute baseline reading, and the fluorescence response at each timepoint was calculated as the change in fluorescence (ΔF = F - F_0_) normalized to the baseline (F_0_). To calculate an integrated cAMP, data points after agonist addition were summed. Each biological replicate represents the average of at least two technical replicates.

### Electrophysiology

Whole cell patch clamp experiments were performed as previously described.^44^ Briefly, HEK 293 µOR stable cells were transfected with 0.7 μg GIRK1-F137S^45^ and 0.1 μg tdTomato (tdT), per well. HEK 293 cells were transfected with GIRK1-F137S, 0.1 μg tdT, 0.35 μg µOR, with or without 0.7 μg GFP-KCTD12, per well. Experiments were performed at least 24 hr after transfection in a high potassium extracellular solution composed of (in mM) 120 KCl, 25 NaCl, 10 HEPES, 2 CaCl_2_, and 1 MgCl_2_ (pH 7.4). Solutions were delivered to a recording chamber using a gravity-driven perfusion system with exchange times of ∼1 s. Cells were voltage-clamped at −60 mV using an Axopatch 200B amplifier (Molecular Devices). Patch pipettes with resistances of 3-7 MΩ were filled with an intracellular solution composed of (in mM): 140 KCl, 10 HEPES, 3 Na_2_ATP, 0.2 Na_2_GTP, 5 EGTA, 3 MgCl_2_ (pH 7.4). Inward GIRK currents were induced with perfusion of 100 nM or 10 µM DAMGO. Desensitization of currents was measured over 60 s DAMGO application. Recordings were analyzed using Clampfit (Molecular Devices) and Prism (GraphPad). Desensitization of the DAMGO-induced currents was calculated as follows: 100 ∗ (1 – (amplitude prior to DAMGO washout) / (peak amplitude following DAMGO application)). The tau of desensitization was calculated from the peak amplitude to DAMGO washout, fit to a single exponential curve.

### Shotgun proteomics data access

RAW data and database search results have been deposited to the ProteomeXchange Consortium via the PRIDE partner repository^46^ with the dataset identifier PXD031415.

### Targeted proteomics data access

Raw data and SRM transition files can be accessed, queried, and downloaded via Panorama^47^ https://panoramaweb.org/MOR-APEX.url.

### Code availability statement

Unbiased mass spectrometry RAW data for APEX proximity labeling experiments were analyzed using the MaxQuant algorithm (version 1.5.5.1) for the identification and quantification of peptides and proteins.^34^ Peptide ion intensities produced by MaxQuant were summarized to protein intensities using the R package MSstats (version 3.23.1).^13^

Targeted proteomics (Selected Reaction Monitoring, SRM) assay generation was performed using Skyline.^38^ The resulting data was analyzed with Skyline for identification and quantification of peptides^38^ and MSstats (version 3.10.6) was used for statistical analysis.^13^

The over-representation analysis (ORA) for Gene Ontology terms was performed using the enricher function from R package clusterProfiler (version 3.99.0).^36^ Cellular location changes were measured using gene set enrichment analysis as implemented in the R package fgsea (v 1.17).^37^

Data independent acquisition RAW mass spectrometry data were analyzed with Spectronaut (Biognosys) using direct DIA analysis default parameters for the identification and quantification of proteins. Peptide ion intensities from Spectronaut were summarized to protein intensities using MSstats (version 3.22.1)^13^ implemented in the R package artMS (version 1.8.3) with default settings.

Affinity purification mass spectrometry (AP-MS) RAW data for EYA4 were searched against the SwissProt Human (downloaded 01/2017) using the default settings for MaxQuant (version 1.6.3.3).^34^ Protein-protein interaction scoring for EYA4 was performed using with both SAINTexpress (version 3.6.3)^39^ and CompPASS^40^. Protein abundances for mutant and wildtype EYA4 AP-MS were compared using MSstats (version 3.14.1).^13^ Protein interaction network was visualized using Cytoscape (version 3.8.1).^41^

Electrophysiology recordings were analyzed using Clampfit (Molecular Devices) and Prism (GraphPad).

## Extended data figures

**Extended Data Figure 1.**
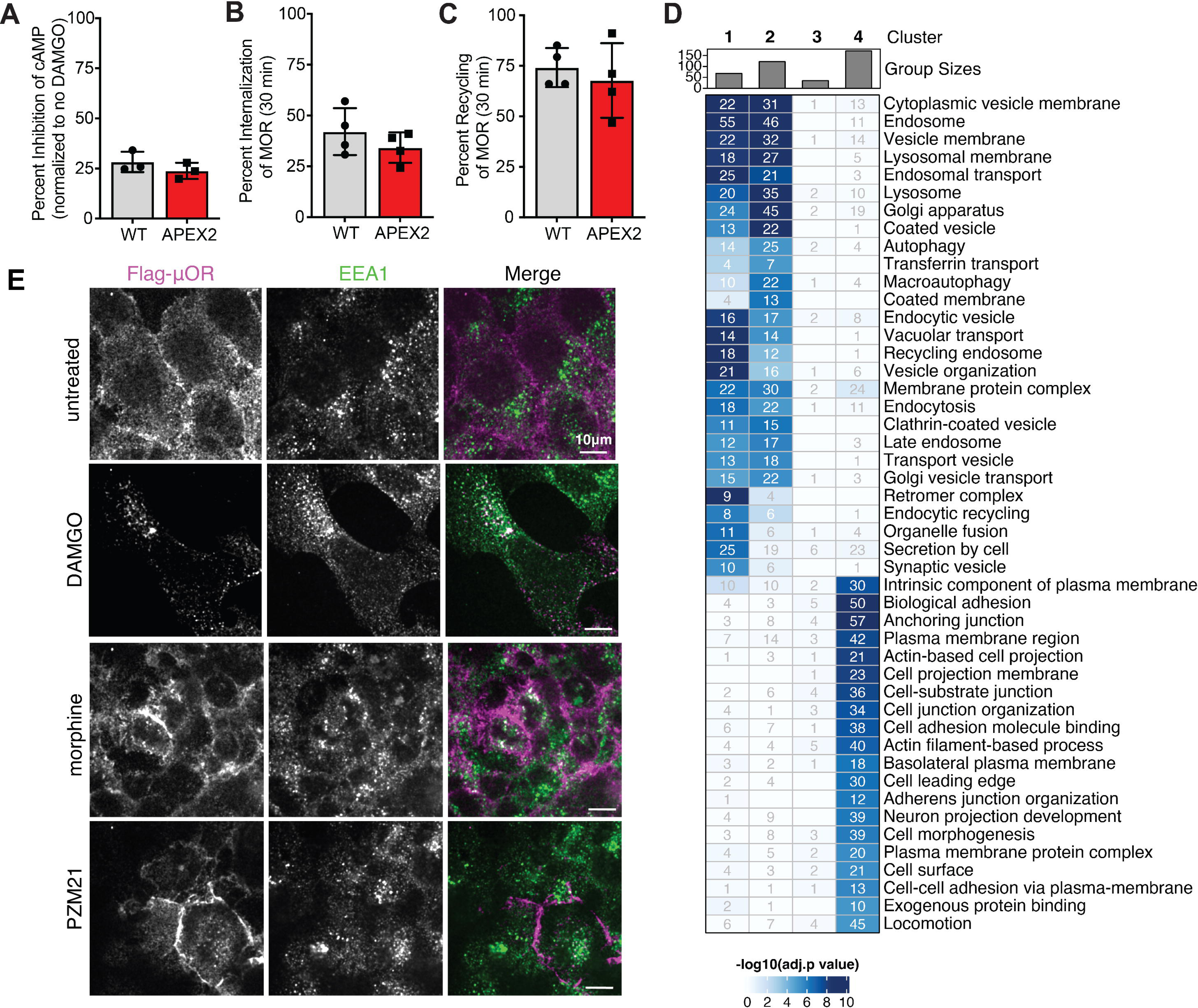
APEX-tagged *μ*OR remains functional and ligand-dependent proximal interaction networks of *μ*OR are enriched for proteins indicating cellular location. A. Receptor signaling was measured using a commercial cAMP biosensor (pGloSensor-20F). cAMP accumulation was measured after ∼10 minutes of DAMGO/isoproterenol incubation and normalized to isoproterenol alone. Data from three independent experiments are presented as mean ± SEM. B. Comparison of agonist-stimulated receptor internalization as assayed by loss of cell surface immunoreactivity and measured by flow cytometry comparing untreated (control) and treated samples (10 μM DAMGO, 30 min). Data from four independent experiments are presented as mean ± SEM. C. Comparison of cell surface recovery of receptors (‘recycling’) following 30 min of DAMGO application (10 μM), agonist removal, and a 30 min recovery period in the presence of antagonist (10 μM). Data from four independent experiments are presented as mean ± SEM. D. The heatmap shows all significantly enriched gene ontology terms (adjusted p-value < 0.05) among the proteins that significantly change in the proximal protein environment of the μOR upon activation with DAMGO, morphine, or PZM21 including the number of proteins that match the gene ontology terms. Cluster 1-4 refer to the clustering of the heatmap in Figure 1B. E. Colocalization of μOR with endosomes to monitor receptor trafficking following activation. HEK293 cells stable expressing the μOR with an N-terminal Flag-tag were activated with 10μM DAMGO, morphine, or PZM21 for 10min. The receptor was imaged using anti-Flag. Endosomes were marked with anti-EEA1. Scale bar is 10μm.

**Extended Data Figure 2.**
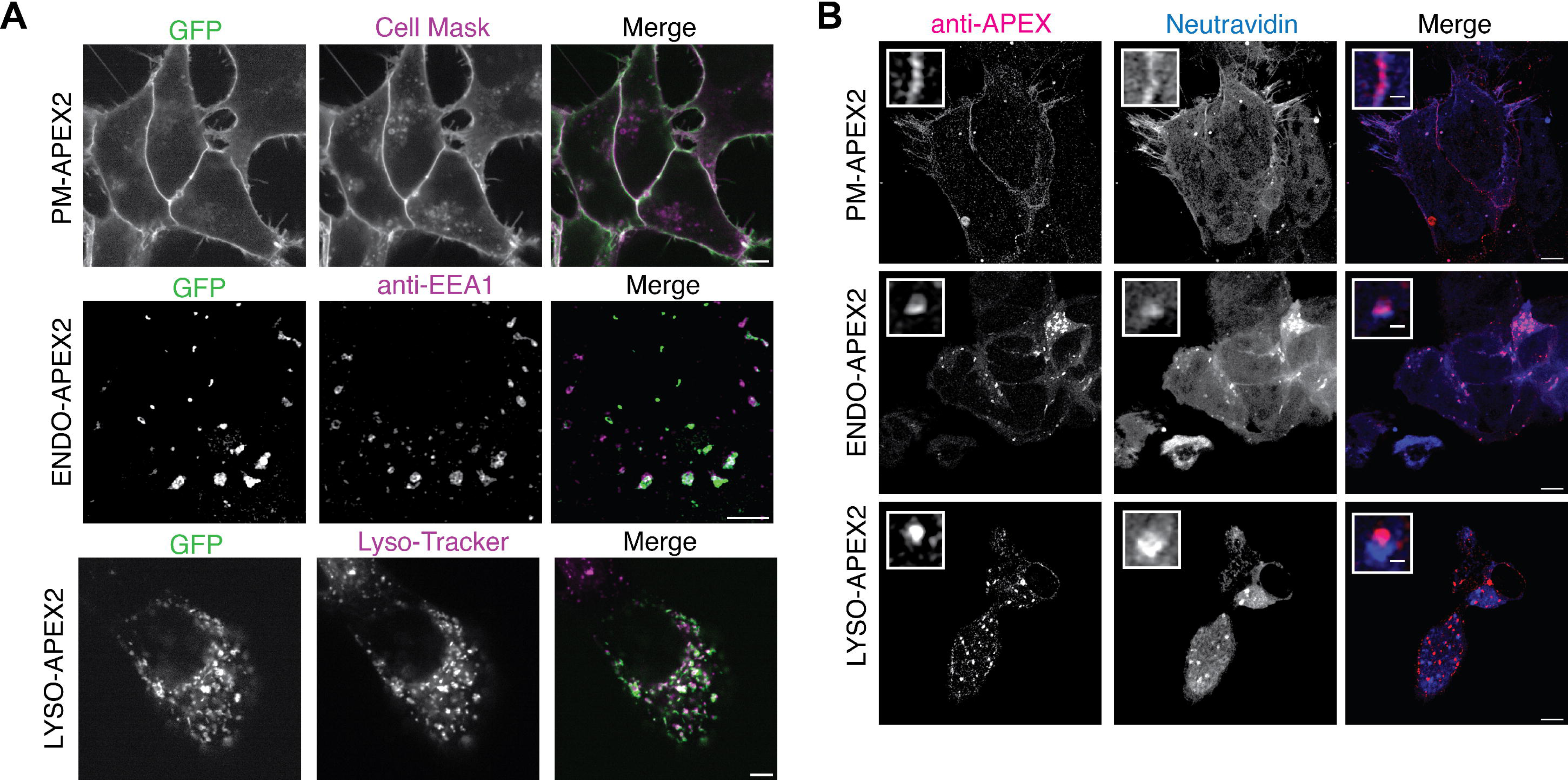
Validation of spatial reference cell lines and *μ*OR trafficking. A. Colocalization of location markers with APEX2 spatial references. HEK293 cells stably expressing PM-APEX2, ENDO-APEX2, or LYSO-APEX2 were imaged with either Cell Mask to mark the plasma membrane, anti-EEA1 to mark endosomes, or Lyso-Tracker to mark lysosomes. N=3 independent biological replicates, representative example shown, scale bar is 5μm. B. Colocalization of biotin with APEX2 spatial reference constructs. Localization of biotin following APEX-mediated proximity labeling was probed with Neutravidin and APEX2 constructs were detected with anti-APEX. N=3 independent biological replicates, representative example shown, scale bar is 5μm.

**Extended Data Figure 3.**
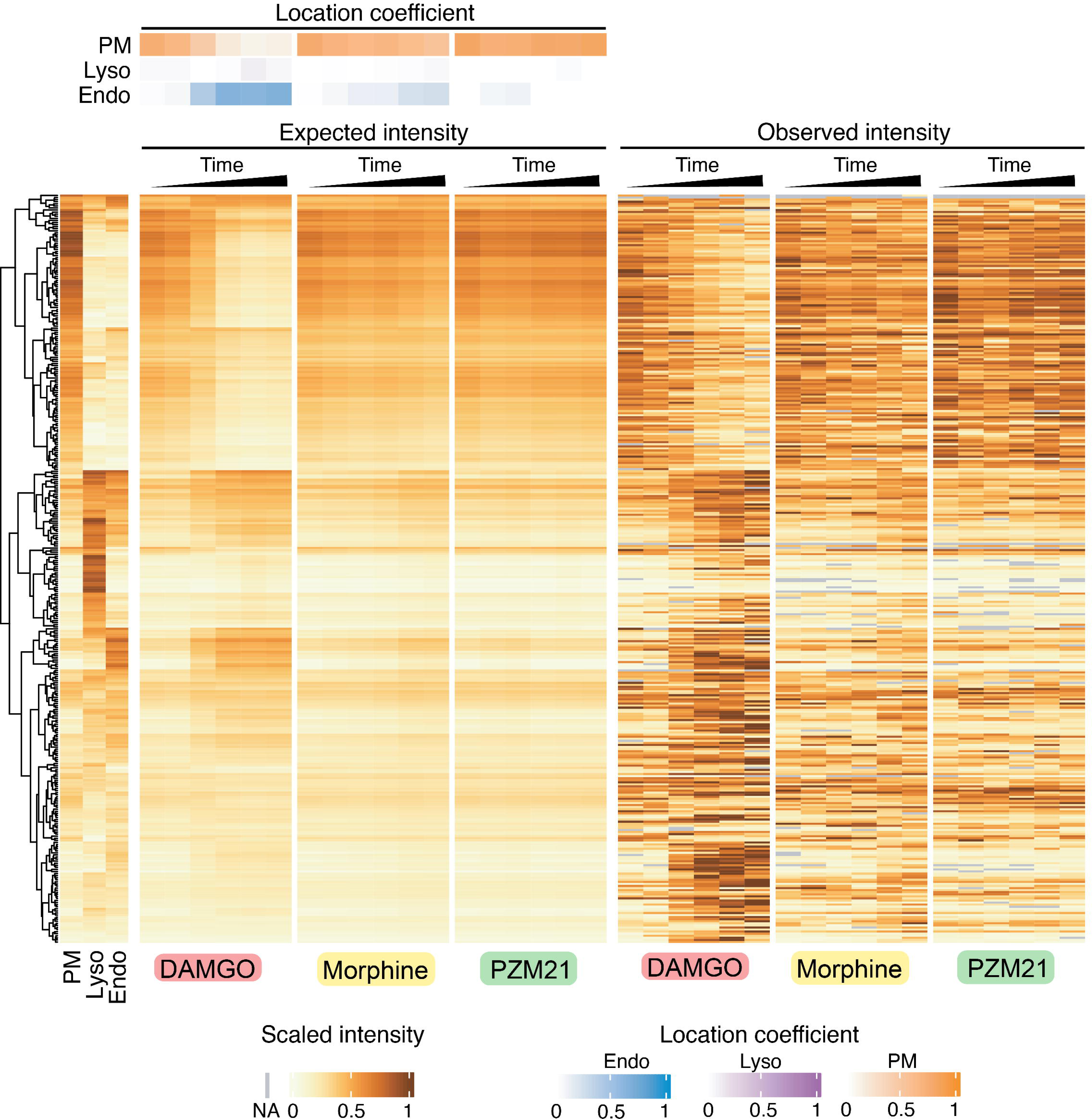
Prediction of expected protein intensities based on location coefficients. The heatmap shows the location specific proteins that were selected by pairwise comparison of the spatial reference data and their scaled intensity measured across the spatial references (left side of the heatmap). Agonist and time point dependent expected protein intensities were estimated by summing the spatial reference protein intensities that were weighted with their respective location coefficient. Observed protein intensities are shown as comparison (right side of the heatmap). Data from three independent experiments are presented as mean.

**Extended Data Figure 4.**
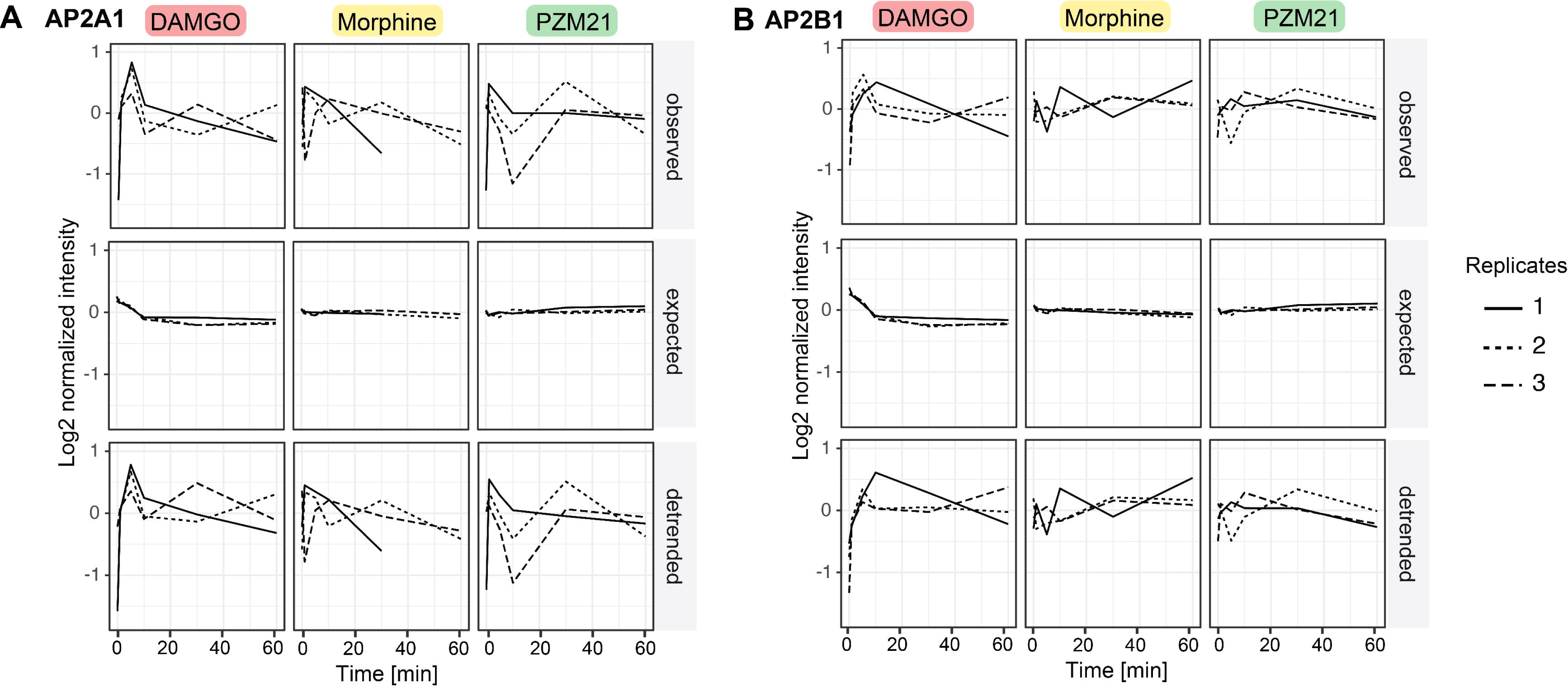
Effect of data detrending for AP2 complex subunits. Data detrending process to dissect localization specific effect from effect of interaction with the receptor for AP2B1 and AP2A1, members of the adaptor protein complex. Three different temporal profiles are depicted for each protein, ligand, and replicate: the initial observed intensities, the expected intensities based on the location specific references, and the intensities after detrending. Data from three independent experiments are presented.

**Extended Data Figure 5.**
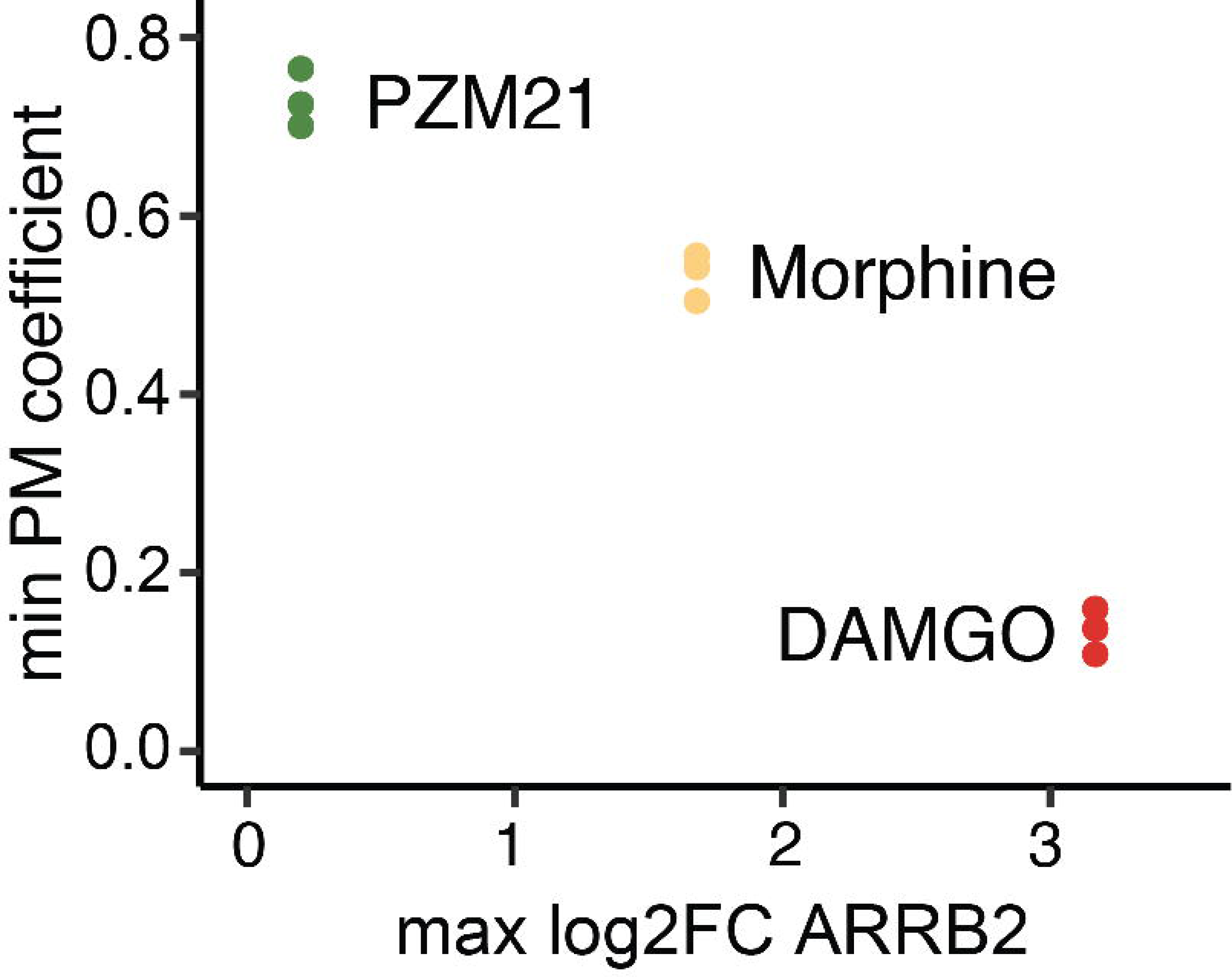
Correlation between ARRB2 engagement upon *μ*OR activation with receptor trafficking. Correlation between the minimum location coefficient calculated for the plasma membrane (PM) and the maximum ARRB2 log2FC over the time course of μOR activation with DAMGO (red), morphine (yellow) and PZM21 (green).

**Extended Data Figure 6.**
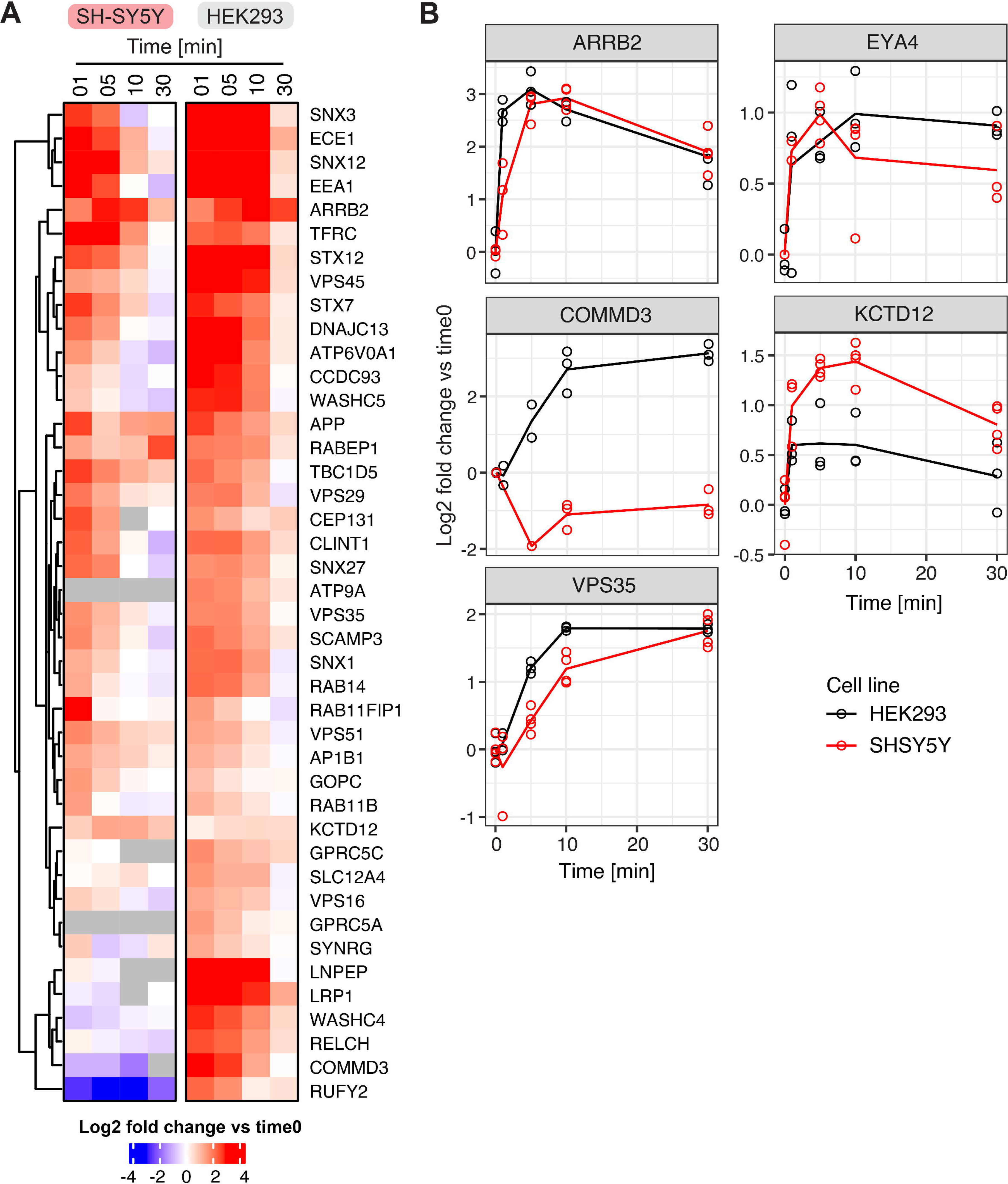
Comparing the DAMGO-dependent proximal proteome changes of the *μ*OR in HEK293 and SH-SY5Y cells. A. Comparison of μOR-APEX experiment upon activation with DAMGO in HEK293 and SH-SY5Y neuroblastoma cells. Heatmap focuses on all proteins significant for DAMGO in HEK293 data depicted in Figure 3A. Data from three biological replicates are presented as mean. B. Temporal profile for selected proximal interactors of the μOR in HEK293 and SH-SY5Y cells. Line charts represent the log_2_ fold change over the time course of receptor activation with DAMGO in HEK293 (black) and SH-SY5Y (red) cells. Data from three biological replicates are presented.

**Extended Data Figure 7.**
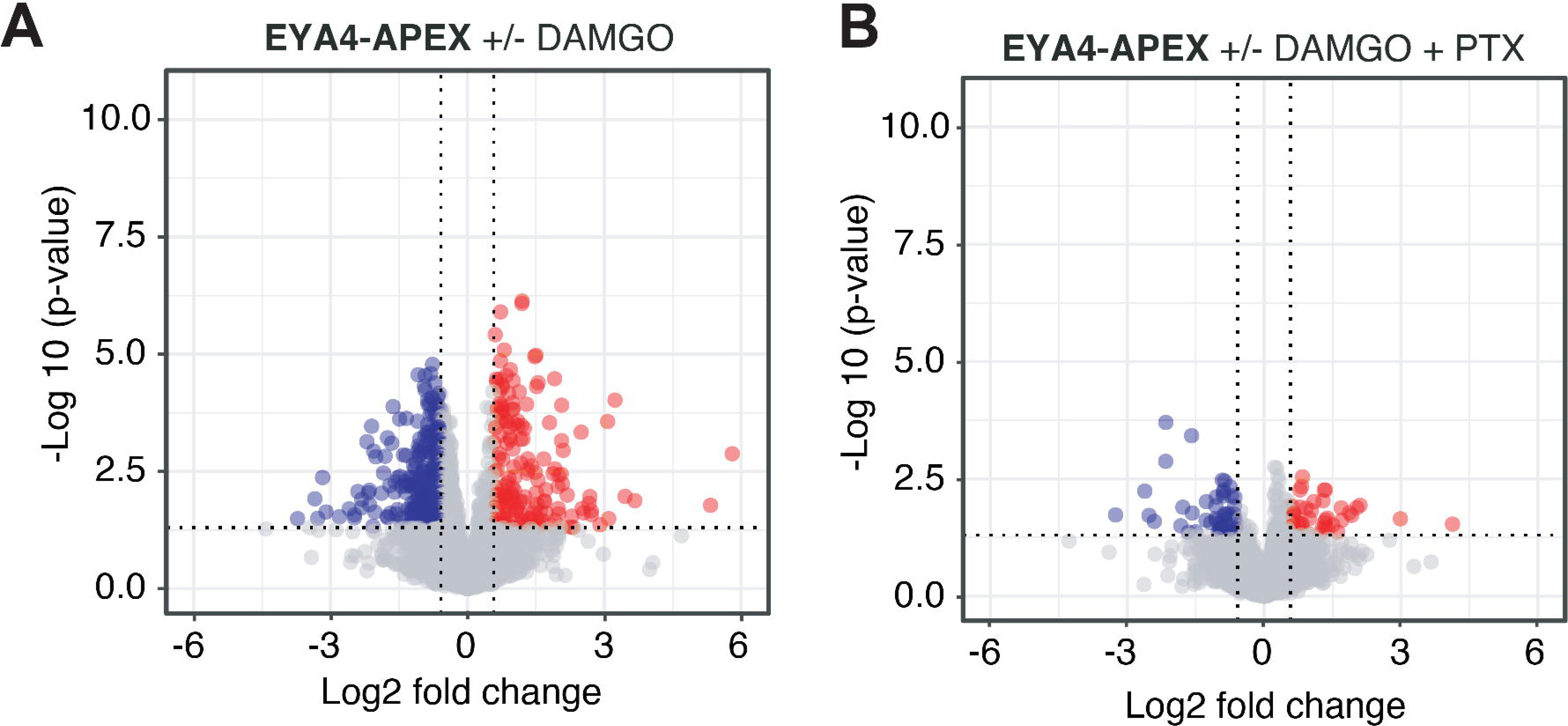
*Changes in proximity labeling of EYA4 upon* μ*OR activation.* Volcano plot depicting log10 p-value and log2 fold change comparing biotin labeled proteins in the proximity of EYA4 in the presence and absence of μOR activation by treatment with 10 μm DAMGO for 10 min and treatment with PTX. Data from three independent replicates are presented as mean.

**Extended Data Figure 8.**
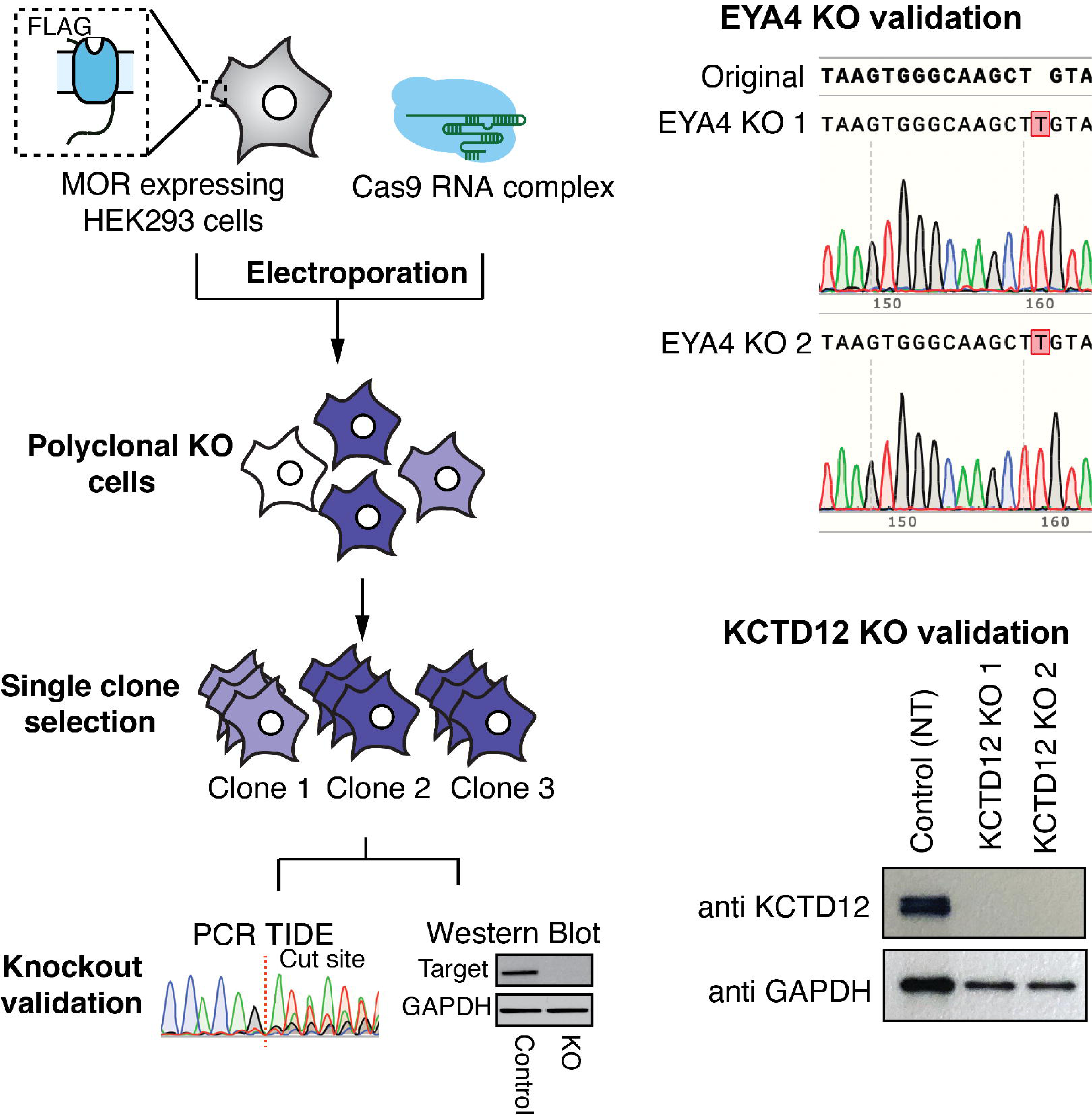
Generation and validation of EYA4 and KCTD12 KO cell lines. KO was validated by PCR and sequencing (EYA4) or Western blot analysis (KCTD12).

**Extended Data Figure 9.**
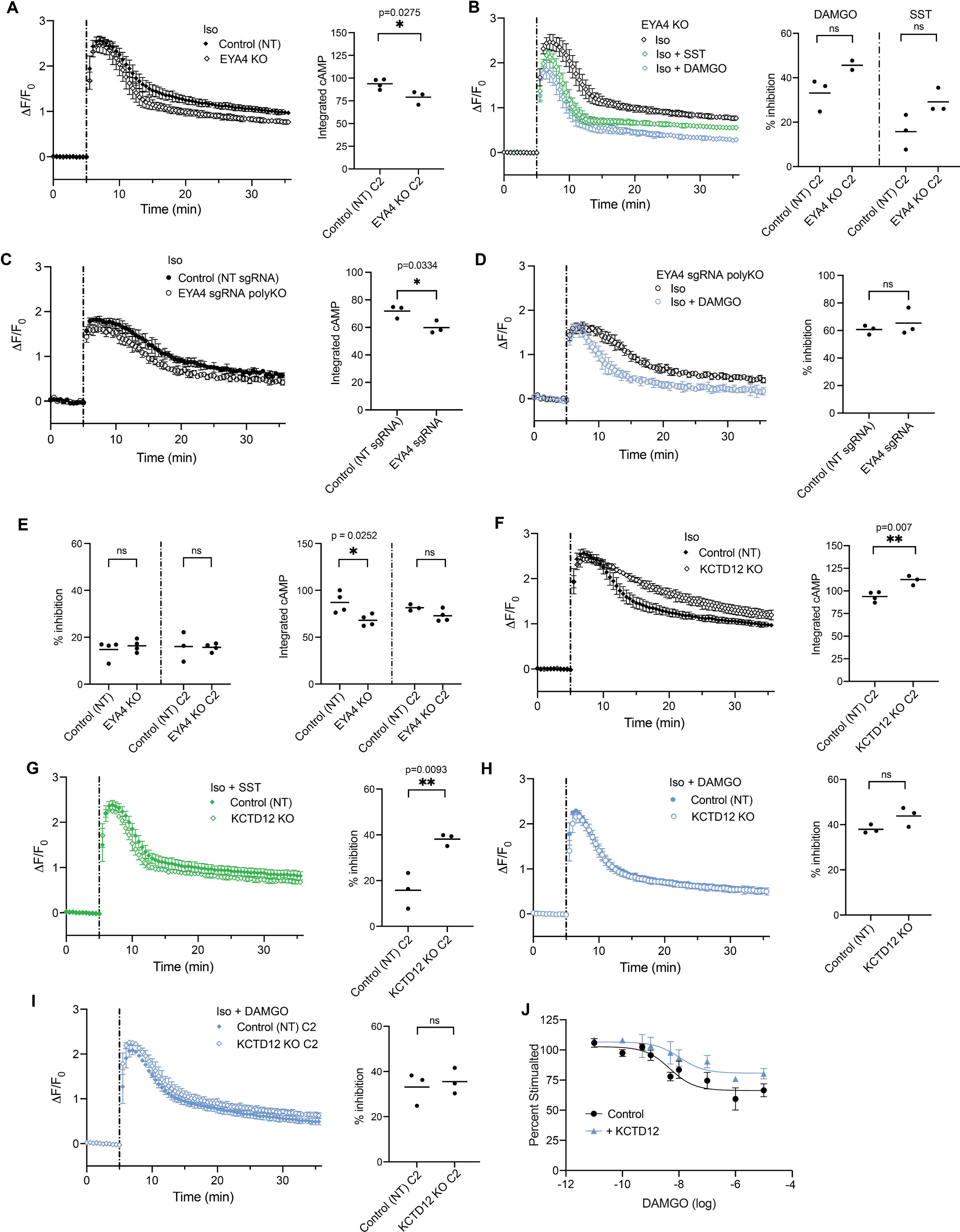
EYA4 and KCTD12 functional validation. A. cAMP activity in control non-targeting (NT) (closed) and EYA4 KO (open) HEK293 cells stably expressing the μOR. Change in fluorescence intensity of cAMP biosensor upon stimulation with 100 nM isoproterenol (Iso) is plotted. B. cAMP activity EYA4 KO cells upon stimulation with Iso without (black) and with co-application of 1 μM somatostatin (SST, green) or 10 μM DAMGO (blue) is plotted. Iso curve is repeated from panel A. C. cAMP activity in control non-targeting (NT) (closed) and EYA4-targeted sgRNA (open) polyclonal HEK293 cells stably expressing μOR. Change in fluorescence intensity of cAMP biosensor upon stimulation with Iso is plotted. D. cAMP activity in polyclonal cells modified with EYA4-targeted sgRNA upon stimulation with Iso without (black) and with co-application of 10 μM DAMGO (blue) is plotted. Iso curve is repeated from panel C. E. Percent inhibition of Iso-stimulated cAMP with co-application of DAMGO (left) and integrated Iso-stimulated cAMP (right) in control non-targeting (NT) and EYA4 KO cell lines stably expressing the μOR pretreated with PTX. F. cAMP activity in control (closed) and KCTD12 KO (open) HEK293 cells stably expressing the μOR upon stimulation with Iso. Control curve is repeated from panel A. G. cAMP activity in control and KCTD12 KO cells upon stimulation with Iso and SST. Percent inhibition data for control is repeated from panel B. H. cAMP activity in control and KCTD12 KO cells upon stimulation with Iso and DAMGO for clones used in main figure (circles) with the main text control curve repeated from Figure 5A. I. cAMP activity in control and KCTD12 KO cells upon stimulation with Iso and DAMGO for clones used in supplemental figures (diamonds). J. cAMP activity in WT cells stably overexpressing μOR-APEX2 and transiently overexpressing KCTD12 or an empty vector control upon stimulation with Iso and DAMGO. For all panels, data represent biological replicates, shown as individual data points or mean ± s.d., and significance was determined by unpaired t-test.

**Extended Data Figure 10.**
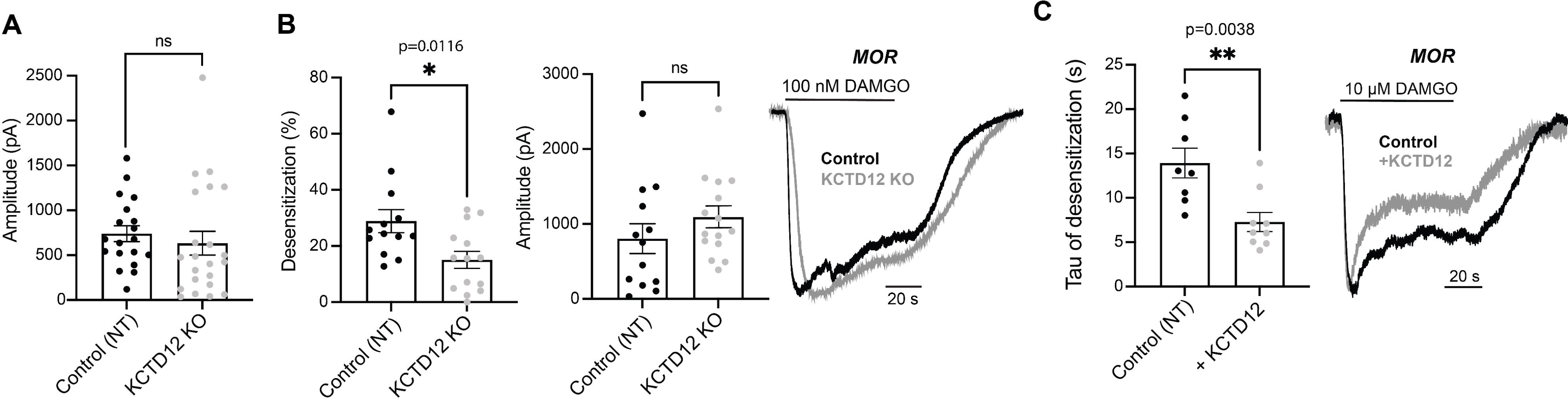
Electrophysiology measurements for KCTD12 KO. A. Summary bar graphs showing the average peak amplitudes of μOR-mediated GIRK currents over 60 s 10 µM DAMGO application, in control and KCTD12 KO HEK cells. Each point represents an individual cell. Error bars represent SEM. B. Left, summary bar graphs showing the average peak amplitudes and percent desensitization of μOR-mediated GIRK currents over 60 s 100 nM DAMGO, in control and KCTD12 KO HEK cells. Each point represents an individual cell. Unpaired t-test, * p=0.0116. Error bars represent SEM. Right, representative whole cell patch clamp recordings of μOR-mediated GIRK currents in response to 60 s 100 nM DAMGO, in control and KCTD12 KO cells. C. Left, Quantification of the tau of desensitization of μOR-mediated GIRK currents over 60 s 10 µM DAMGO application, without (control) and with KCTD12 overexpression in HEK 293T cells. Each point represents an individual cell. Unpaired t-test, ** p=0.0038. Error bars represent SEM. Right, representative whole cell patch clamp recordings showing GIRK currents mediated by μOR activation over 60 s 10 µM DAMGO.

## Supplemental Tables

***Supplemental Table 1*.** The table lists all proteins quantified across the μOR-APEX samples in HEK293 cells and the results of their statistical analysis over the time course of activation with DAMGO, morphine and PZM21.

***Supplemental Table 2.*** Location specific proteins that were quantified by targeted proteomics in the μOR-APEX samples.

***Supplemental Table 3.*** Selection of indicative proteins for subcellular location of the receptor from the pairwise comparison of proteins quantified across the spatial references for the plasma membrane (PM-APEX), early endosome (Endo-APEX), and lysosome (Lyso-APEX).

***Supplemental Table 4.*** The table lists all proteins quantified across the μOR-APEX samples and the results of their statistical analysis over the time course of activation with DAMGO, morphine and PZM21 after detrending of the dataset for location specific effects.

***Supplemental Table 5.*** Ligand-dependent proximal interaction network of the μOR. The table contains all proteins that show a significant fold change for at least one ligand before and after detrending of the data for location-specific effects.

***Supplemental Table 6*.** The table lists all proteins quantified across the μOR-APEX samples in SH-SY5Y cells and the results of their statistical analysis over the time course of activation with DAMGO.

***Supplemental Table* 7.** Targeted proteomics (SRM) assays used to quantify selected proteins in the μOR-APEX dataset including localization markers and potential interactors of the μOR.

